# Harnessing β-Lactam Antibiotics for Illumination of the Activity of Penicillin-Binding Proteins in *Bacillus subtilis*

**DOI:** 10.1101/2019.12.19.881714

**Authors:** Shabnam Sharifzadeh, Felix Dempwolff, Daniel B. Kearns, Erin E. Carlson

## Abstract

Selective chemical probes enable individual investigation of penicillin-binding proteins (PBPs) and provide critical information about their enzymatic activity with spatial and temporal resolution. To identify scaffolds for novel probes to study peptidoglycan biosynthesis in *Bacillus subtilis*, we evaluated the PBP inhibition profiles of 21 β-lactam antibiotics from different structural subclasses using a fluorescence-based assay. Most compounds readily labeled PBP1, PBP2a, PBP2b or PBP4. Almost all penicillin scaffolds were co-selective for all or combinations of PBP2a, 2b and 4. Cephalosporins, on the other hand, possessed the lowest IC_50_ values for PBP1 alone or along with PBP4 (ceftriaxone, cefoxitin), 2b (cefotaxime) or 2a, 2b and 4 (cephalothin). Overall, five selective inhibitors for PBP1 (aztreonam, faropenem, piperacillin, cefuroxime and cefsulodin), one selective inhibitor for PBP5 (6-aminopenicillanic acid) and various co-selective inhibitors for other PBPs in *B. subtilis* were discovered. Surprisingly, carbapenems strongly inhibited PBP3, formerly shown to have low affinity for β-lactams and speculated to be involved in resistance in *B. subtilis*. To investigate the specific roles of PBP3, we developed activity-based probes based on the meropenem core and utilized them to monitor the activity of PBP3 in living cells. We showed that PBP3 activity localizes as patches in single cells and concentrates as a ring at the septum and the division site during the cell growth cycle. Our activity-based approach enabled spatial resolution of the transpeptidation activity of individual PBPs in this model microorganism, which was not possible with previous chemical and biological approaches.

## INTRODUCTION

Bacterial cell shape and integrity are dictated by the rigid external layer that surrounds the cell, known as the cell wall. A major component of this protective coating, which is unique to prokaryotes, is the peptidoglycan (PG), a heteropolymeric structure composed of saccharide backbones that are cross-linked by peptide subunits.(1) PG composition and structure are linked to cell wall thickness and integrity, hence determining sensitivity to cell wall-targeting antibiotics and host immunity.(2) Biosynthesis of the PG requires the coordinated action of multiple bacterial enzymes. Of great significance are the penicillin-binding proteins (PBPs), which mediate the final steps of PG assembly, through polymerization and crosslinking of the PG chains.(3, 4) PBPs are classified based on their molecular weight and conserved amino acid motifs. PBPs are divided into two main subclasses based on their molecular mass; high molecular mass (HMM) and low molecular mass (LMM). HMM PBPs are composed of multiple domains and are involved in the polymerization and crosslinking of the PG chains. Based on the number of reactions that each enzyme can catalyze, HMM PBPs are further classified. While class A proteins are bifunctional and catalyze both polymerization and crosslinking processes, class B PBPs only catalyze the latter. LMM PBPs, which are also referred to as class C, catalyze a different process called D,D-carboxypeptidation, during which the stem pentapeptide is hydrolyzed to a tripeptide.(3, 4) PBPs were discovered and named for their affinity to bind to β-lactam antibiotics such as penicillin, which comprise the largest class of clinically-utilized antibiotics today. Most bacteria possess a relatively large suite of these proteins, between ~4-16 homologs.(1, 4–6) Although the importance of the PBPs as antibacterial targets has long been appreciated, the specific roles of individual PBPs, beyond their catalytic function, has remained elusive due to the lack of appropriate chemical and biological tools.(7, 8) Functional redundancy within the PBPs has compounded the difficulties in establishing the role of individual PBPs in peptidoglycan biosynthesis.(9–11)

Single PBPs have been studied by conjugation to fluorescently labeled protein tags.(10, 12) However, these data alone are insufficient to fully characterize their roles as protein localization does not provide information about the activity state of the PBPs. Moreover, protein tags are bulky and can perturb protein abundance, localization, or function.(13) Alternatively, fluorescent D-amino acids (FDAAs) have been developed that can be incorporated into nascent cell wall, reporting on the formation of new PG material, but do not provide protein homolog-specific information.(14–17) As such, we have pursued the identification of small molecule scaffolds with high affinity for individual homologs that enable selective targeting of individual PBP catalytic activities.

The earliest strategy for detection of PBP activity was tagging of these proteins with radiolabeled penicillin, followed by separation by sodium dodecylsulfate polyacrylamide gel electrophoresis (SDS-PAGE). As suicide inhibitors of the PBPs, radioactive penicillins provide information about the catalytic state of the PBPs, and as such, they have been used in kinetic and inhibitor binding assays.(18–20) More recently, fluorescent probes replaced their radioactive counterparts due to their added advantages including: application to live cells, imaging potential and faster, lower hazard procedures. Bocillin-FL (Boc-FL), a fluorescent derivative of penicillin V is a global probe that typically labels all PBPs in a given organism.(21, 22) Following this advancement, we and others have utilized fluorescence-based assay for profiling PBP inhibition using Boc-FL as the readout probe from live cells.(22, 23) Moreover, fluorescent penicillin-based probes have enabled monitoring of PBP transpeptidation activity in live cells.(21, 24, 25) Previously, we generated fluorescent Ceph C-based probes that demonstrated selectivity for a subset of PBPs and used them to label active PBPs in Gram-positive bacteria.(7) More recently, we developed a class of PBP-selective probes using a β-lactone scaffold and visualized the activity of individual transpeptidases in *S. pneumoniae*.(8) Given the proven success of our probe toolbox, we aimed to develop probes for individual PBPs in *Bacillus subtilis*.

*B. subtilis* is a sporulating Gram-positive organism with a large number of PBPs. In 1972, Strominger demonstrated the presence of five different penicillin-binding components in *B. subtilis*.(26) Today, 16 genes are known to encode PBPs in this microorganism, which include four class A, six class B and six class C PBPs.(8, 9, 12, 27) Some of these PBPs play roles in cell growth and division,(9, 12, 28) while others are involved in synthesis of the spore PG.(29–33) However, as described previously, the specific tasks of individual members have been difficult to assign due to functional redundancy of these enzymes.(9, 10, 12, 32) In order to facilitate the development of chemical tools to study cell growth and division in *B. subtilis*, we assessed the PBP inhibition profile of a library of β-lactam antibiotics in this model microorganism using the fluorescent whole cell assay mentioned above. Briefly, *B. subtilis* cells were treated with increasing concentrations of different β-lactam antibiotics, followed by Boc-FL as the readout probe to determine which PBPs were blocked by the antibiotic. Subsequently, cells were lysed and the membrane fraction was analyzed by SDS-PAGE. As expected, several PBP targeting profiles were observed for each β-lactam structural subclass, among which the high affinity of carbapenems for PBP3, a transpeptidase known for low sensitivity to β-lactams, stood out. In order to investigate the activity of PBP3, we designed and synthesized probes based upon meropenem. Our meropenem-based probes retained the PBP binding profile of the core meropenem molecule and specifically targeted PBP3 and PBP5. We then applied the probes to the *B. subtilis* DK654 strain, in which *dacA*, encoding PBP5, is knocked out to enable specific visualization of PBP3. Super resolution fluorescent imaging revealed a distinct localization pattern for PBP3, with a high concentration of activity at the septal region and the separation site in the mid-to late-divisional stage. Interestingly, inhibition of the only essential PBP in *B. subtilis*, PBP2b, via pretreatment of the cells with mecillinam did not change the labeling pattern, which suggests that PBP3 acts independently and complements PBP2b activity.

## RESULTS

### Selectivity profile of β-lactams for *B. subtilis* PBPs

Herein, twenty-one β-lactam antibiotics from different subclasses (penicillin, cephalosporin, carbapenem, penem and monobactam) were tested in *B. subtilis PY79* strain to determine their PBP binding profile (**Fig. S1**). This is the first example of a comprehensive *in vivo* profiling of the PBP activity in this model microorganism. Results are presented as gel images and graphs in **Figures S2**. For each gel, the fluorescence intensity of individual bands was measured and compared with the corresponding PBP band from a parallel sample not treated with antibiotics. The average of these relative intensity values from two independent assays was plotted against antibiotic concentration to calculate the IC_50_ values of individual PBPs for each compound (Table 1). A compound was considered selective for a specific PBP if the measured IC_50_ was at least 4-fold lower than that of the next most inhibited PBP (smaller IC_50_). When the IC_50_ value for a second, third or fourth PBP fell below this threshold, that compound was considered co-selective for all of the most inhibited PBPs. Selectivity criteria were the same as described previously.(22, 23)

**Table 1.** IC_50_ and MICs of β-lactams in *B. subtilis* PY79.

| β-lactam | MIC<br>(μg/ml) | IC<sub>50</sub>(s)<sup>a</sup> (μg/ml) | | | | | | selectivity |
| --- | --- | --- | --- | --- | --- | --- | --- | --- |
|  |  | PBP1a/1b | PBP2a | PBP2b/H | PBP3 | PBP4 | PBP5 |  |
| Aztreonam | 16 | <0.01 | >1000 | 26.13±20 | 192.7±95.3 | 5.8±0.2 | >1000 | 1 |
| Faropenem | 0.125 | 0.00059±0.00008 | 0.011±0.006 | 0.0051±0.0018 | 0.94±0.11 | 0.009±0.005 | 0.20±0.01 | 1 |
| Doripenem | 0.0625 | >1000 | <1000 | <1000 | 0.12±0.01 | >1000 | 0.23±0.03 | 3, 5 |
| Meropenem | 0.0625 | >1000 | <1000 | <1000 | 0.26±0.01 | ≥1000 | 0.22±0.01 | 3, 5 |
| Mecillinam | 16 | >1000 | 6.72±2.68 | 1.07±0.23 | 89.6±41.3 | >1000 | >1000 | 2a, 2b |
| Penicillin V | 32 | *<0.01 | 0.00042±0.0003 | <0.01 | 2.04±1.08 | <0.01 | 0.62±0.30 | 2a, 2b, 4 |
| Penicillin G | 16 | > 1000 | 0.0043±0.0005 | 0.019±0.001 | 1.4±0.2 | 0.0090±0.0013 | 0.62±0.35 | 2a, 2b, 4 |
| Ampicillin | 8 | *<0.01 | 0.0036±0.003 | 0.0094±0.0037 | 0.042±0.025 | 0.0037±0.003 | 0.12±0.04 | 2a, 2b, 4 |
| Amoxicillin | 4 | *0.0010±0.0002 | 0.074±0.018 | 0.088±0.007 | 0.14±0.04 | 0.076±0.023 | 1.11±0.05 | 2a, 2b, 4 |
| Piperacillin | 64 | 0.0012±0.006 | 0.16±0.10 | < 1.0 | 1.9±0.8 | 0.030±0.003 | 11.3±1.6 | 1 |
| Methicillin | 0.25 | >1000 | < 0.01 | < 0.1 | 9.0±0.8 | 0.11±0.03 | 148.8±43.5 | 2a, 2b |
| Oxacillin | 0.25 | >1000 | 0.056±0.026 | 0.061±0.006 | 66.8±9.8 | <0.1 | >1000 | 2a, 2b, 4 |
| Cloxacillin | 0.125 | >1000 | 0.09±0.03 | 0.13±0.06 | <1000 | 0.022±0.002 | >1000 | 2a, 2b, 4 |
| 6-APA | 8-256 | >1000 | 110.6±55.1 | >1000 | >1000 | >1000 | 4.1±0.2 | 5 |
| Cephalexin | 0.5 | >1000 | <0.1 | <0.1 | 24.1±4.8 | 254.4±73.1 | >1000 | 2a, 2b |
| Cefuroxime | 0.125 | 0.0018±0.0006 | 27.9±3.8 | 0.08±0.04 | 22.2±1.0 | 0.015±0.003 | >1000 | 1 |
| Cefotaxime | 0.0625 | 0.0013±0.0001 | <100 | <0.1 | 0.99±0.07 | <0.1 | >1000 | 1, 2b, 4 |
| Ceftriaxone | 0.25 | 0.0075±0.0007 | 16.3±0.2 | 0.066±0.025 | 2.6±0.06 | 0.020±0.006 | >1000 | 1, 4 |
| Cephalothin | 0.03125 | 0.0026±0.0011 | 0.0065±0.004 | 0.0076±0.0029 | 11.8±2.2 | 0.006±0.003 | 108.1±3.7 | 1, 2a, 2b, 4 |
| Cefsulodin | 8 | 0.0098±0.0003 | 38.0±20.6 | 0.66±0.09 | 138.6±13.2 | 0.12±0.02 | >1000 | 1 |
| Cefoxitin | 1 | 0.014±0.001 | 0.72±0.23 | 0.19±0.007 | 1.04±0.06 | 0.014±0.003 | 0.19±0.01 | 1, 4 |

<sup>a</sup> IC<sub>50</sub>s were determined as described in Materials and Methods, based on 10-fold concentration steps, using a log(inhibitor)-versus-response-variable slope (four parameters) and least square (ordinary) fit with GraphPad Prism software. Average of two individual experiments are provided. An IC<sub>50</sub> of >1,000 was assigned when GraphPad Prism reported an ambiguous number and significant inhibition was not seen at 1,000 μg/ml. In cases where GraphPad Prism resulted in ambiguous values, IC<sub>50</sub> was reported as lower than the concentration that caused > 50% inhibition of that PBP.
\*Asterisk indicates that the examined β-lactam resulted in 50% inhibition of that PBP, but complete inhibition was not observed up to 1000 $\mu\text{g/ml}$ .

Five of the tested β-lactams, including aztreonam, faropenem, piperacillin, cefuroxime and cefsulodin were selective for PBP1, which is a class A HMW PBP in this microorganism. 6-aminopenicillanic acid (6-APA) selectively inhibited PBP5, the only LMM PBP active in the vegetative state of *B. subtilis*, at sub-micromolar concentrations. The finding that 6-APA acted as PBP5 inhibitor is in contrast with our previous work that indicated the low affinity of this compound for any PBPs in *S. pneumoniae* and *E. coli*.(22) Aside from PBP1 and PBP5, co-selective inhibition of multiple PBPs was observed by the remainder of the β-lactams (Table 1).

### Multiple compounds displayed high affinity for class A PBPs

All tested cephalosporins, except cephalexin, were selective or co-selective inhibitors of PBP1, which is one of the class A PBPs in *B. subtilis*. Aztreonam and faropenem, a monobactam and a penem, also showed high affinity for this PBP. On the other hand, piperacillin was the only compound from the penicillin subclass that resulted in complete inhibition of this PBP within the tested concentrations (Fig. 1; **Fig. S2**). These findings were in contrast with a previous report on titration of *B. subtilis* membrane lysates by [^14^C]penicillin G followed by SDS-PAGE and fluorography, which revealed that PBP1a, 1b and 2c saturate at very low concentrations, followed by PBPs 2a and 4.(34) Moreover, another report suggested penicillins such as mecillinam, methicillin and ampicillin as strong inhibitors of both PBP1 and 2 from *B. subtilis* membrane lysates.(35) In another study on *Bacillus megaterium*, which is similar to *B. subtilis*, whole cells were treated with [^14^C]penicillin G and PBP1 was suggested as the main target for this β-lactam.(36) The results of these studies are inconsistent with our work, which indicates the potentially significant difference between the activity of PBPs *in vivo* versus in cell lysates, demonstrating the utility of our strategy.(22, 32)

**Figure 1.**
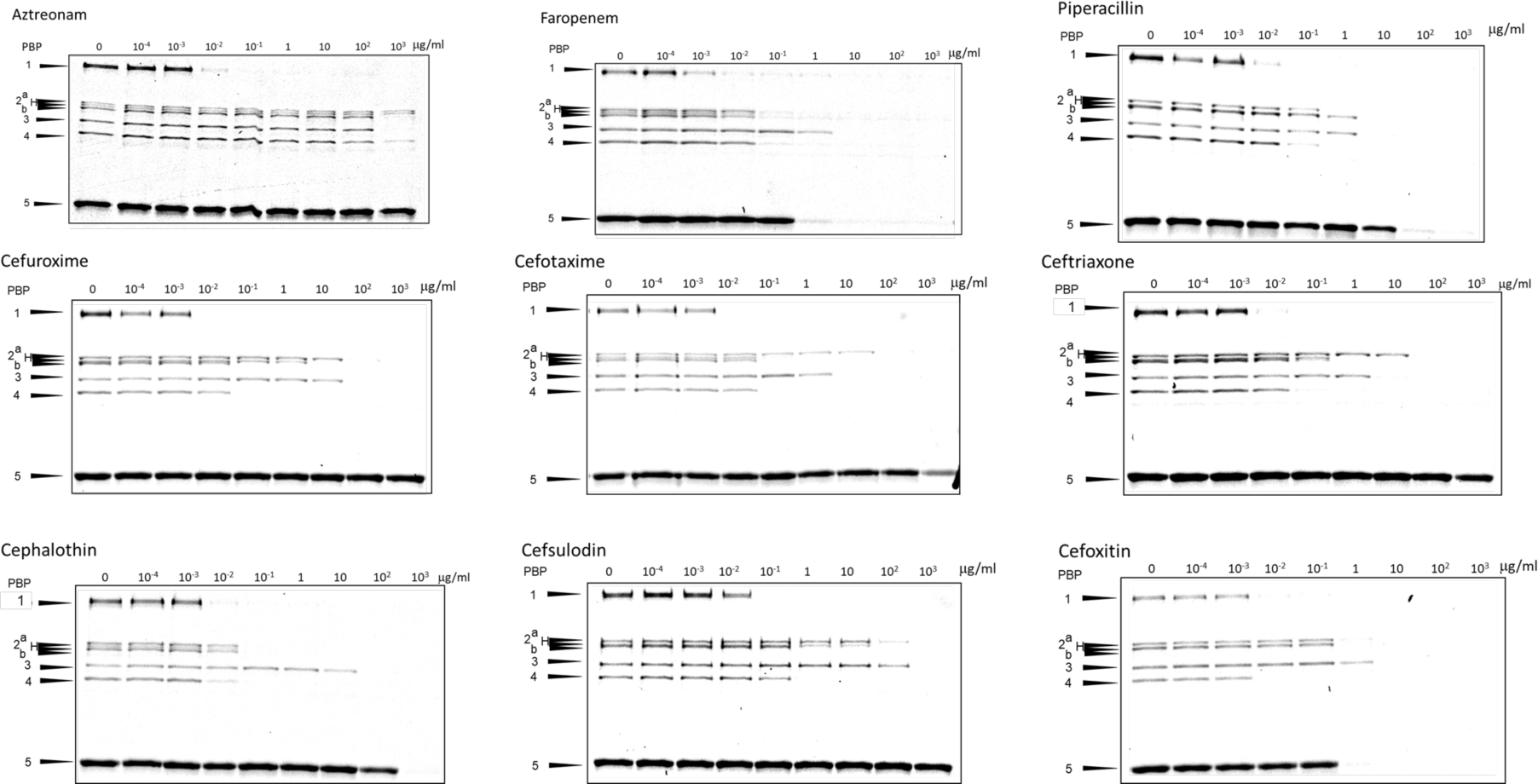
Complete inhibition of PBP1 by aztreonam, faropenem, piperacillin and cephalosporin compounds. Representative SDS-PAGE gel image for titration of *B. subtilis* PY79 cells by these antibiotics over a defined concentration range. Gel band quantitation for the presented gels, with the standard deviations of the results from two independent experiments are plotted in **Figure S2**.

### Penicillins are potent inhibitors of PBP2a and 2b

Based on the molecular weight and labeling experiments with PBP knock out strains, we determined that PBPH is co-migrating with PBP2b under our assay conditions (**Fig. S3**). PBPH is a class B transpeptidase with high structural similarity to PBP2a.(9) Deletion of *pbpH* does not cause phenotypic changes. However, Δ*pbpA*Δ*pbpH* double mutants are not viable.(9) Comparison of the fluorescence intensity between wild-type and the Δ*pbpH* strain labeled by Boc-FL, a measure of TP activity, revealed no significant difference in the relative activity for most of the PBPs (**Fig. S3** and **Table S1**). Moreover, titrations with ampicillin (PBP2a, 2b and 4 inhibitor) and methicillin (PBP2a and 2b inhibitor), were performed with DK694 (Δ*pbpH*) cells (**Fig. S4**) and gel bands for all PBPs were quantitated to confirm the relatively minor contribution of PBPH to the IC_50_ values for the co-migrating PBP2b. IC_50_ values for individual PBPs were comparable, and within 2-fold in most cases (**Table S2**). These results are consistent with the fact that PBPH activity is minimal in early log phase and becomes prominent only in late exponential phase. Given the minimal contribution of PBPH activity to fluorescent labeling observed in the PBP2 region, PBP2b and PBPH were integrated together, and the resulting value was used as a measure of PBP2b activity.

In this study, all tested penicillins were potent inhibitors of PBP2a and PBP2b, with calculated IC_50_ values in the submicromolar range for most compounds, with the exception of 6-APA as mentioned previously. Mecillinam exclusively targeted PBP2a and PBP2b over the tested range of concentrations (Fig. 2, **S2**). Mecillinam is a specific inhibitor of PBP2 in *E. coli*, preventing lateral cell wall elongation, causing formation of spherical cells incapable of dividing, ultimately leading to cell death.(37–39) PBP2a has long been believed to play roles in maintaining rod shape in *B. subtilis* and mutation in PBP2a has been found to cause twisted cells.(9, 40, 41) PBP2b, encoded by *pbpB*, is the only PBP that is essential for growth in *B. subtilis*, which is attributed to its major role in septum formation.(10, 41, 42) More recently, it was revealed that the transpeptidation activity of PBP2b is dispensable and cells containing deactivated PBP2b maintain normal growth and morphology.(10, 11) Thus, the development of chemical tools that can selectively target the activity of these transpeptidases can complement the genetic studies and deepen our knowledge about these PBPs, which will be the subject of future work.

**Figure 2.**
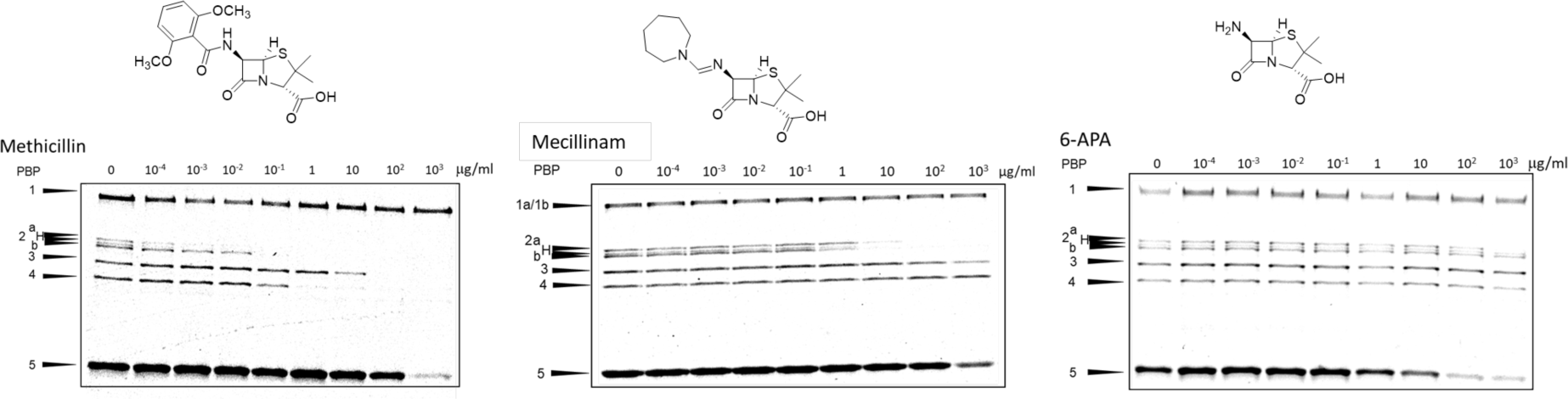
Representative SDS-PAGE gel image for inhibition of the PBPs by methicillin (left), mecillinam (center) and (+)-6-aminopenicillanic acid (right) in *B. subtilis* over a range of antibiotic concentrations. Whole cells were treated with various concentrations of antibiotics and subsequently labeled with 5 μg/ml Boc-FL. Most penicillins were potent inhibitors of PBP2a and PBP2b, as exemplified by methicillin. Mecillinam exclusively inhibited these two PBPs over the tested concentration range. 6-APA is a weak inhibitor of PBP5 activity.

### Carbapenems displayed a distinct inhibition profile

All tested compounds showed the highest affinity for class A HMW PBPs, except for 6-APA (PBP5 inhibitor) and the two tested carbapenems; doripenem and meropenem, which selectively inhibited PBP3 and PBP5, a class B transpeptidase and a D,D-carboxypeptidase, respectively (Fig. 3A). Interestingly, these two carbapenems yielded very low MIC values, 0.0625 μg/mL, at which most PBPs are only minimally inhibited (Fig. 3A and Fig. 4). To investigate whether growth in the presence of meropenem might affect the PBP activity profile after an extended period of exposure, we grew *B. subtilis* cells with a sub-MIC concentration of meropenem (0.01 μg/mL, 2 h) and examined the PBP activity profile. No significant inhibition of any PBP occurred under these conditions, perhaps suggesting an additional role for meropenem that leads to cell death (**Fig. S5**).

**Figure 3.**
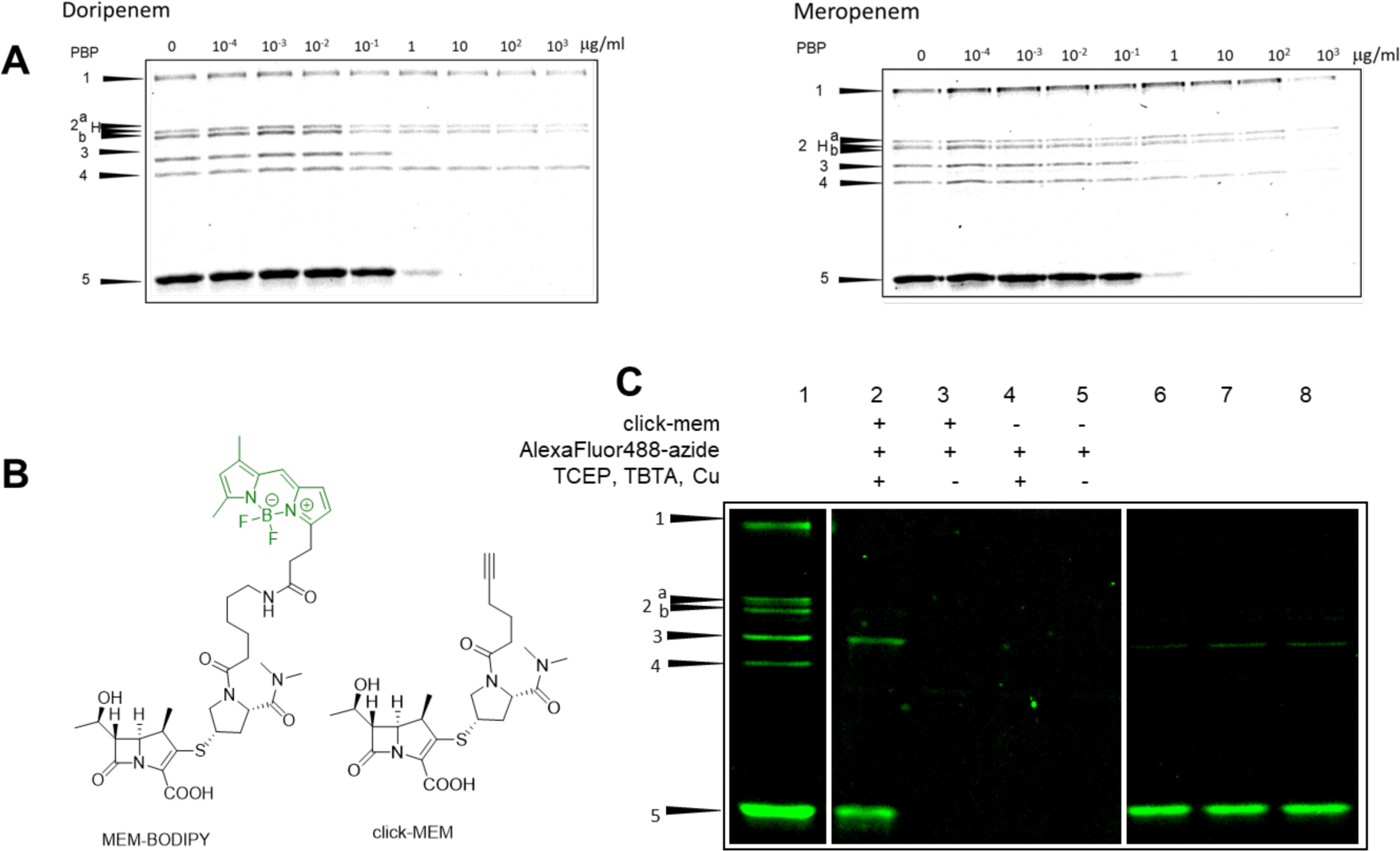
Probes to target PBP3 and PBP5 in *B. subtilis.* **A**) Doripenem and meropenem selectively target PBPs 3 and 5. **B**) Chemical structure of fluorescent (MEM-BODIPY) and clickable meropenem analogs (click-MEM); **C**) PBP labeling profile of meropenem-based probes in *B. subtilis* cells assessed by SDS-PAGE. **1**: 10 µM bocillin-FL (control) **2-5**: treatment by 5 µM click-MEM (2,3) or no probe (4,5) followed by reaction with different combinations of click reagents. Only lane 2, where cells were treated with the probe and reacted with AlexaFluor488-azide in the presence of all click reagents resulted in fluorescent labeling of PBPs. **6-8**: treatment with 2 µM MEM-BODIPY for 10 min (6), 20 min (7) or 30 min (8). Labeling intensity plateaus after 20 minutes.

**Figure 4.**
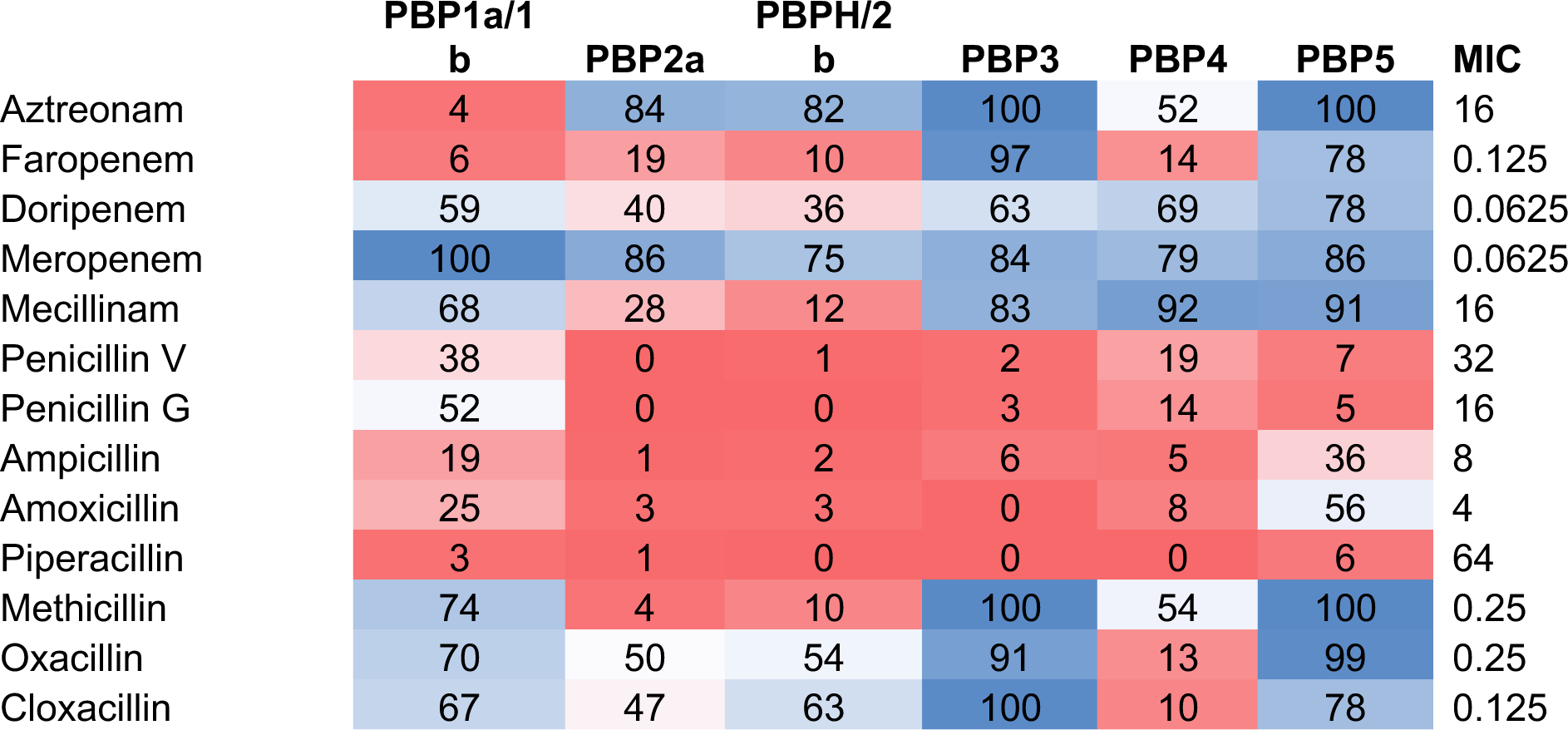
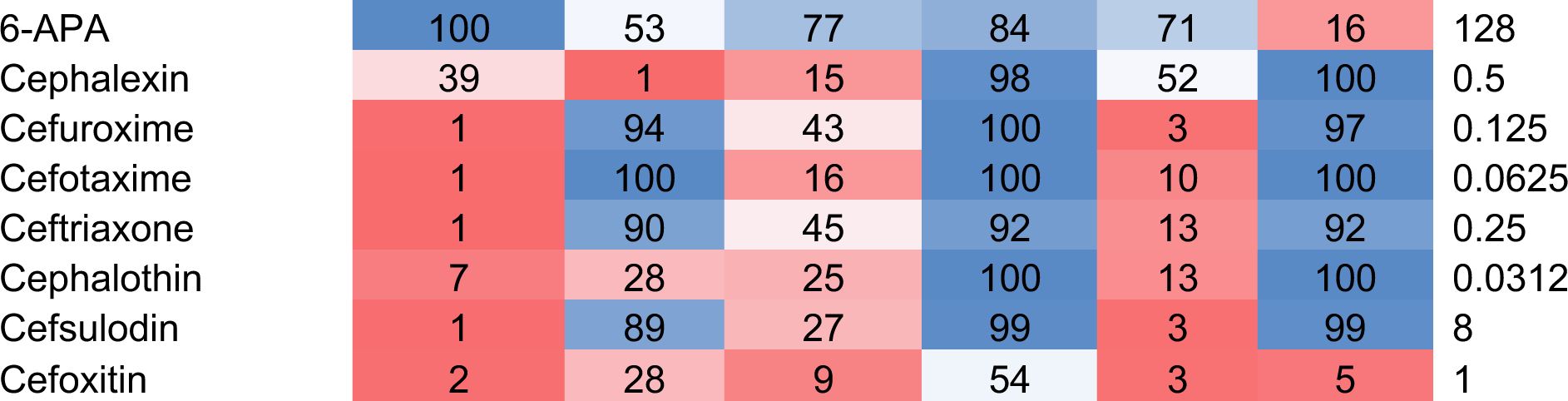
PBP inhibition at MIC illustrated as a heat map to summarize the level of PBP inhibition at growth inhibitory concentrations. The numbers in each cell is the average of % Boc-FL labeling of each PBP at MIC concentration, from two separate experiments (i.e., low numbers equate to more protein inhibition). Since titrations were done in 10-fold intervals, the Boc-FL labeling at the concentration closest to MIC was used. Low Boc-FL labeling (red) demonstrates high inhibition of that PBP at that concentration, whereas blue indicates high levels of Boc-FL labeling/low PBP inhibition.

### Lethal target(s) of β-lactams in *B. subtilis*

Relying on our titration and growth inhibition results, we sought to determine the main killing target(s) for the tested β-lactams. Minimum inhibitory concentration (MIC) values were plotted against IC_50_ values for every PBP separately (**Fig. S6**). No conclusive relationship could be drawn between inhibition of any specific PBP and cell growth. This is in contrast to our previous work in *S. pneumoniae* that yielded an obvious relationship between the MIC value and inhibition of one of the essential PBPs, PBP2x.(43) Given the essentiality of PBP2b, one might expect that substantial inhibition (>50%) of this protein would be required for cell killing. This is not the case with aztreonam, meropenem, 6-APA, oxacillin or cloxacillin, further supporting the recent demonstration that abolishment of its TP activity is tolerated.(10, 11) However, a closer look at the profile of the PBPs targeted at inhibitory concentrations of individual compounds reveals some interesting trends within different β-lactams subclasses. Penicillins completely blocked PBP2a and PBP2b, as well as one or more additional PBPs at sub-MIC levels (**Table S3**). The only exception is 6-APA, which consists of only the penicillin core with no side chain. At growth inhibitory concentrations, 6-APA failed to significantly block any PBPs. All cephalosporins, excluding cephalexin, targeted PBP1 under sub-MIC concentrations. The same results were observed with aztreonam and faropenem, a monobactam and a penem, respectively (**Table S3**, Fig. 4). While this cumulative data might implicate all these PBPs as potential lethal targets in *B. subtilis*, our results with 6-APA, as mentioned, along with meropenem and doripenem, immediately contradict this hypothesis. Considering the different growth conditions for PBP labeling (i.e., PBS) *vs* MIC assays (i.e., LB), it is possible that individual PBPs may have varying roles and importance during different stages of cell division.

### Development of meropenem-based probes to target PBP3

According to our results, among the transpeptidases in *B. subtilis*, PBP3 has the lowest susceptibility to β-lactams. The only exceptions were the carbapenems (Table 1). Given the unique affinity of the carbapenems for PBP3 (Fig. 3A), we designed chemical probes based upon meropenem to target the activity of this protein. Although none of the tested inhibitors only targeted PBP3, we pursed the development of a meropenem-based probe as this scaffold should label PBP3 and PBP5 and could be used in conjunction with a *pbp5* deletion mutant (∆*pbpC*; DK695) to readily visualize PBP3 activity, as we have done in previous studies. (8, 25)

The secondary amino group located in the pyrrolidine ring of meropenem was used to attach a fluorophore to enable visualization of the TP activity (**MEM-BODIPY**; **Scheme S1**). Another analog containing an alkyne side chain, **click-MEM**, was produced to enable installment of different reporter and affinity tags through the copper(I)-catalyzed Hüisgen azide-alkyne cycloaddition (CuAAc; Fig. 3B). Both probes retained specificity for PBP3 and PBP5, identical to the core meropenem molecule (Fig. 3C). Assessment of the PBP targeting profile in strains in which PBP3 or PBP5 were knocked out, DK654 (*∆pbpC*) and DK695 (*∆dacA*), respectively, confirmed these proteins as the targets of our meropenem-based probes (**Fig. S7**). To further confirm this, we tagged the *B. subtilis* proteome with click-MEM and installed a biotin affinity tag via CuAAc. Biotinylated proteins were subsequently enriched by Neutravidin-agarose beads, subjected to proteolytic digestion by trypsin and the resulting peptide mixture was analyzed by LC-MS/MS. The obtained data were searched against the *B. subtilis* FASTA amino acid sequence database and confirmed that PBP3 and PBP5 were enriched as expected (**Fig. S9**).

### PBP imaging in *B. subtilis* using β-lactam probes

PBPs of *B. subtilis* have been previously visualized *in vivo* using fluorescent protein tags or antibodies.(10, 12) We were interested in using our meropenem-based probes to visualize the activity of PBPs and establish their potential for imaging studies. We utilized the MEM-BODIPY FL probe to monitor PBP3 activity in *B. subtilis DK654*, in which PBP5 is knocked out and therefore, our MEM-BODIPY FL probe specifically targets PBP3. The images showed that PBP3 localized in the septal region as a ring in early-to-mid divisional cells and moved to the separation site during late divisional stages (Fig. 5). We also observed either patch-like or polar distribution of fluorescence signal in single cells (Fig. 5, **S10**). Fluorescent signal was significantly reduced upon pretreatment of the cells with 10 μg/mL meropenem, which blocks PBP3 activity (**Fig. S11** and **S12**). This is in contrast with a previous study that used fluorescent protein tags to localize PBPs in *B. subtilis* and reported that a fluorescent PBP3 construct distributed around the cell periphery and was not enriched at the site of the septum.(12) This difference in results could be due to the fact that the chemical labeling indicates the location of PBP3 activity, but our findings are consistent with a recent study in which *B. subtilis* PBP3 was immunolabeled and showed predominant localization at mid-cell and the cell poles.(10) We conclude that either the fusion of the fluorescent protein to PBP3 caused localization artifacts or PBP3 is dynamic in localization and activity.

**Figure 5.**
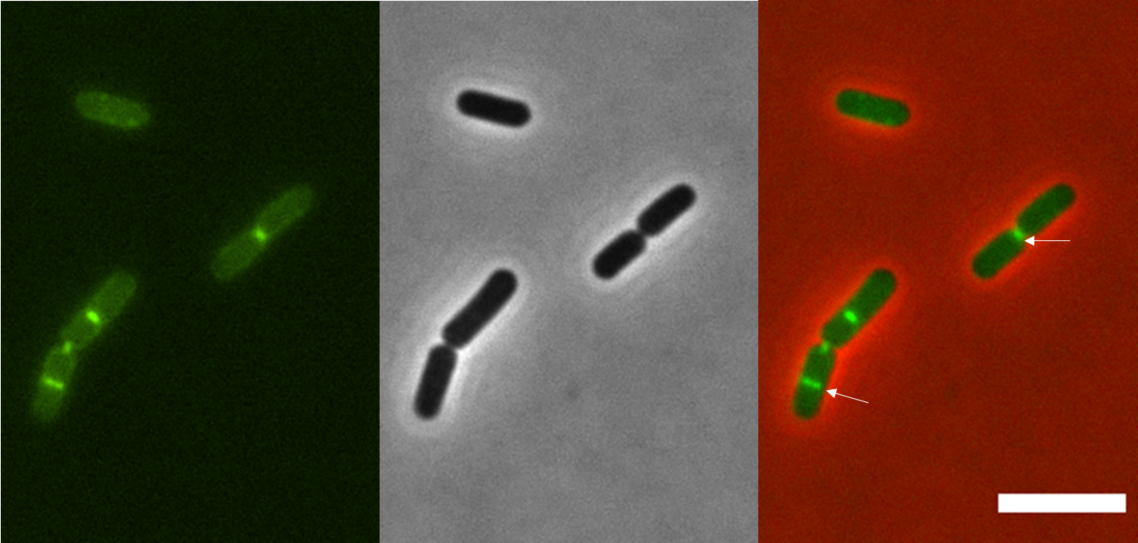
Visualization of PBP3 activity in live *B. subtilis DK654* cells (*∆dacA*). Exponentially growing *B. subtilis DK654* cells were labeled with 2 μM MEM-BODIPY FL and imaged at 100× magnification. Fluorescent image is shown on *left*, phase contrast in *middle* and both overlaid on *righ*t. PBP3 activity localizes to septal region in early-to mid-divisional stages, and concentrates at the division site later on (demonstrated by the white arrows). Scale bar = 5 μm.

Next, we sought to employ our Click-MEM probe for the same purpose, since the alkyne functionality would enable installation of different reporter groups in the future. Performing the labeling and click chemistry in phosphate buffer saline led to lysis of the majority of the cells. We found that performing these steps in minimal media would overcome this problem. However, the background fluorescence signal remained high in comparison to the MEM-BODIPY FL probe (**Fig. S10**).

PBP2b is the only essential PBP in *B. subtilis*, with established roles in cell division. It was recently shown that the transpeptidation activity of PBP2b is dispensable and PBP3 becomes the essential transpeptidase once PBP2b is deactivated.(10) In order to target PBP3 in the absence of PBP2b activity, DK654 cells were pretreated with 10 μg/mL mecillinam, which we have shown to selectively inhibit PBP2b (Table 1, **Fig. S2**). Fluorescence imaging of cells pretreated with mecillinam revealed a similar labeling at the septum of the dividing cells and patch-like labeling of the single cells when compared to cells that were not pretreated, and thus had full PBP2b catalytic activity (**Fig. S13**).

## DISCUSSION

Although PBP TP activity is fundamental to normal bacterial physiology, it has not been possible to conclusively determine which PBPs may be most critical for TP activity in live cells. PBPs in *B. subtilis* have long been studied as the targets of β-lactam antibiotics. The covalent nature of the complex formed between β-lactams and PBPs has made them invaluable tools, aside from their therapeutic utility, for both isolation(26) and analysis of this family of enzymes.(7, 21, 22, 24, 33, 35, 44) Studies of β-lactam selectivity in *B. subtilis* and other microorganisms have been conducted for two main purposes: 1) β-lactam binding studies to identify those PBPs directly related to measured MICs, which reveals that those PBPs are potentially essential for the growth of microorganism; and 2) studies on strains resistant to specific β-lactams to identify which PBPs show a change in sensitivity for that antibiotic and if those PBPs play a role in resistance.(6) Radioactive penicillins were initially exploited to tag PBP activity in bacterial lysates in such experiments.(22, 26, 33–36, 40, 45–47) In spite of the invaluable information that these *in vitro* studies have provided about the affinity of β-lactams for PBPs, it must be noted that membrane preparation could potentially affect and change protein activity.(48) Therefore, we performed a comprehensive study and titrated *B. subtilis* cells with an extensive library of β-lactams, using a fluorescent whole cell assay.

*B. subtilis* has been extensively studied as a Gram-positive model microorganism. As part of its cell wall biosynthesis machinery, *B. subtilis* possesses four class A PBPs that polymerize PG, known as PBP1, 4, 2c and 2d, encoded by genes *ponA*, *pbpD*, *pbpF* and *pbpG*, respectively.(32, 49–51) Construction of mutant strains lacking one or multiple class A PBP-encoding genes revealed that only loss of PBP1 results in significant phenotypic and PG structure changes.(28, 52, 53) Simultaneous deletion of PBP1 and PBP4 accentuates PG structural defects with a profound decrease in growth rate.(52) PBP2c and PBP2d are known to play roles mainly in spore PG synthesis.(32) In our titration studies, we found many selective inhibitors for PBP1, which could be used as selective inhibitors to further investigate the roles and regulation of this PBP. Additionally, several penicillins and a few cephalosporin compounds including ceftriaxone, cephalothin and cefoxitin, resulted in very low IC_50_ values for PBP4. Overall, all tested compounds except carbapenems, mecillinam and 6-aminopenicillanic acid, favored PBP1, PBP4 or both, as their main target in *B. subtilis*. This substantial preference for class A HMM PBPs is in contrast with our preceding results in other microorganisms, in which LMM PBPs were favored. In our previous work on *S. pneumoniae*, as another Gram-positive microorganism, the most inhibited PBPs were 3, followed by 2x, which are LMM and class B HMM PBPs, respectively.(22) Similarly in *E. coli*, as a rod shape model organism, we found that PBP3, a class B transpeptidase, as well as PBPs 4, 7 and 8, all LMM PBPs, were targeted more frequently compared with class A HMM PBPs.(22)

The main lethal target(s) of β-lactams in *B. subtilis* is still under debate. Two main strategies have been employed to investigate lethal targets of β-lactams in different microorganisms. One method is to identify the PBPs that are inhibited the most at concentrations leading to growth inhibition.(35) An alternative method is to determine which PBPs show altered activity in mutant strains, assuming that decreased affinity of certain PBPs has caused resistance.(33) Early gel-based studies on *B. subtilis* performed prior to resolution of PBP2a and PBP2b, identified PBP2 as the main killing target of β-lactams in this microorganism.(33, 40, 45) PBP2 was identified as the main killing target of cloxacillin in *B. subtilis* strain Porton and carbenicillin-resistant mutants.(33, 45) However, affinity of PBP2 from mutant strains for penicillin did not change in this study, which could raise the possibility of distinct modes of interaction for different β-lactam molecules with the penicillin-binding site of the enzyme.(33) Upon resolving the PBP2 band into PBP2a and PBP2b, the former was suggested as the lethal target(45), although PBP2b was never completely eliminated as the other potential option.(12) Moreover, *B. subtilis* strains lacking PBP2a or PBP3 have been shown to become more sensitive to β-lactams.(10) We did not find any specific correlations between the inhibition of any PBP and inhibition of growth in *B. subtilis* (**Fig. S6**). However, we noted that in general, at the MICs, at least one HMM PBP must be fully blocked (Fig. 4).

Among the currently available β-lactam antibiotics, carbapenems have gained a unique status due to their broad spectrum of antibacterial activity and resistance to hydrolysis by different classes of β-lactamases or even inhibition of these enzymes in some cases.(54, 55) Previous *in vitro* studies have shown that carbapenems induce morphological changes in *Pseudomonas aeruginosa* strains in a time- and concentration-dependent manner.(56, 57) Meropenem and doripenem are particularly potent for the inhibition of PBP2 and PBP3 in *P. aeruginosa* and PBP2 in *E. coli*, both rod-shaped Gram-negative organisms.(58) Morphological studies in *P. aeruginosa* revealed that both meropenem and doripenem cause filamentation, with a central oval swelling, and no significant cell wall breakage and lysis.(57, 59, 60) In our titration studies, both doripenem and meropenem showed a distinct PBP binding profile, selectively inhibiting PBPs 3 and 5. PBP3, encoded by *pbpC*, is a class B HMW transpeptidase which is mostly expressed during the vegetative phase. The activity of PBP3 is dispensable in wild type cells. Mutations in *pbpC* were shown to cause helical cells, with no significant effect on peptidoglycan composition(10, 11, 40) and increased susceptibility to β-lactams.(10, 40) Analysis of the primary sequence of PBP3 revealed significant similarity to *Enterococcus faecium* PBP5, *S. aureus* PBP2a and *Escherichia coli* PBP2.(61) More recently, the essential role of PBP3 in the absence of PBP2b activity in cell division was discovered.(10, 11) Visualization of PBP3 activity in live *B. subtilis*, by chemical probes as presented in this work, also suggested a potential role for this PBP in cell division, which seems to proceed independently from PBP2b TP activity. Future studies will benefit from this methodology to further explore PBP3 and PG transpeptidation in *B. subtilis* and other bacterial systems. Finally, biotin-mediated enrichment of meropenem-tagged proteins from whole cells enabled the isolation of PBP3 and PBP5, highlighting the utility of our chemical tools for investigation of the PBPs and the proteins associated with them.

## CONCLUSION

Herein, we presented the first extensive analysis of PBP inhibition profiles of the β-lactam antibiotics in *B. subtilis* cells. Several compounds showed dose-dependent selectivity for a single PBP, while the remainder of the molecules inhibited multiple PBPs. Additionally, mecillinam stood out as a specific inhibitor of PBP2b (predominantly) and PBP2a (to a lesser extent). Meropenem and doripenem resulted in a distinct binding profile, exclusively inhibiting PBP3 and PBP5. Based on these results, chemical probes were designed and synthesized using the meropenem core to target PBP3. These probes enabled examination of PBP3 activity in live *B. subtilis* cells, which mainly localized as a ring during early divisional stages and moved to the central septal region and the division site during later stages. This finding is unprecedented and suggests a potential role for PBP3 during the division of *B. subtilis* cells. Overall, this work highlights the significance of PBP-selective chemical tools to complement the existing molecular and cell biology tools, in order to augment our understanding of bacterial growth and division processes.

## MATERIALS AND METHODS

β-Lactam titration and detection of PBPs, synthesis and characterization of probes, microscopy and additional experimental protocols are discussed in detail in the Supporting Information.

## ACKNOWLEDGEMENTS

This work was supported by the National Institutes of Health (R01 GM128439-01A1 to E.E.C. and R35 GM131783 to D.B.K.), a Sloan Research Fellow Award (E.E.C.), a University of Minnesota Interdisciplinary Doctoral Fellowship (S.S.), and the University of Minnesota Department of Chemistry. The authors thank Y. Zhao and P. Villalta for assistance with the operation mass spectrometers. Mass spectrometry was carried out in the Analytical Biochemistry Shared Resource of the Masonic Cancer Center, University of Minnesota, supported in part by the Cancer Center Support Grant NIH CA077598.

## SUPPORTING INFORMATION

Supporting information is available and contains additional gel and microscopy images, data tables, proteomics and MIC data, materials and methods, and small molecule synthesis information and characterization data.

## TOC Graphic

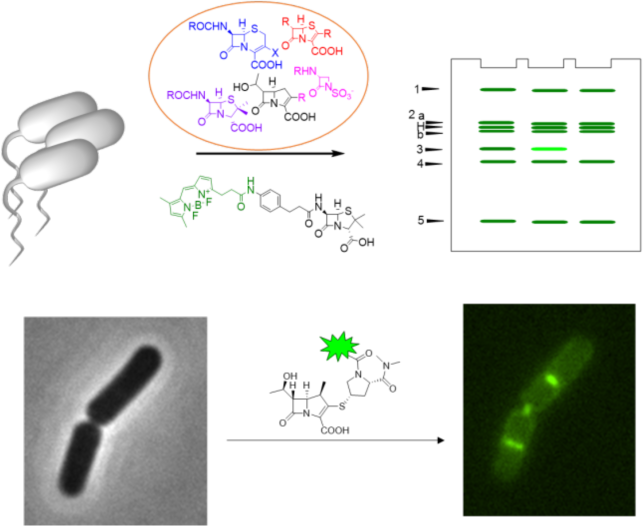

