## Supplementary material for "Harnessing β-Lactam Antibiotics for Illumination of the Activity of Penicillin-Binding Proteins in *Bacillus subtilis*": SI file

**Supplemental Tables:**

**Table S1.** Relative transpeptidase activity of different PBPs in *B. subtilis* PY79 (wild type) *vs* DK694 (Δ*pbpH*).

**Table S2.** $IC_{50}$ determination of ampicillin and methicillin in *B. subtilis* PY79 (wild type) *vs* DK694 (Δ*pbpH*).

**Table S3.** PBP inhibition profile shown by β-lactams at sub-MIC concentrations.

**Table S4.** Inhibition of Boc-FL labeling of PBPs in *B. subtilis* PY79.

**Supplemental Figures:**

**Figure S1.** Structures of the β-lactam antibiotics used in this study.

**Figure S2.** Representative SDS-PAGE gels (one of two independent experiments) and the representative graphs for β-lactam titration of the PBPs in *B. subtilis* PY79.

**Figure S3.** PBP activity profile of *Bacillus subtilis* PY79 (wild type) and mutant strains.

**Figure S4.** Representative SDS-PAGE gels for β-lactam titration of PBPs in *B. subtilis* DK694 (Δ*pbpH*).

**Figure S5**. PBP activity profile of *B. subtilis PY79* cells grown in the presence of 0.01 μg/mL meropenem.

**Figure S6.** Graphs of MIC *vs* $IC_{50}$ for PBPs in *B. subtilis*.

**Figure S7.** PBP labeling profile of meropenem-BODIPY FL in *B. subtilis 3610* (wild type), *DK654* (Δ*dacA*) and *DK695* (Δ*pbpC*).

**Figure S8.** Meropenem targets in membrane and cytosolic fractions of *B. subtilis 3610* strain.

**Figure S9.** Volcano plot of proteins enriched by click-MEM probe.

**Figure S10.** Visualization of PBP3 activity in live *B. subtilis DK654* cells.

**Figure S11.** Meropenem pretreatment inhibits PBP labeling by MEM- probes, gel-based analysis.

**Figure S12.** Pretreatment with meropenem removes PBP3 labeling by MEM-BODIPY FL probe, microscopy imaging.

**Figure S13.** Visualization of PBP3 activity in live *B. subtilis DK654* cells pretreated with mecillinam.

**Scheme S1.** General synthesis scheme for meropenem-based probes.

**Additional Protocols and Data**

General Materials and Methods

Bacterial Culture

β-Lactam titration and detection of PBPs

Probe Concentration Determination

Probe Labeling

Gel-Based Analysis

LC-MS/MS Analysis of Meropenem Targets

Antimicrobial Susceptibility Assay

Fluorescent Microscopy

Probe Synthesis and Characterization

NMR spectra

% Activity of Total

| PBP | PY79 | DK694 |
|---|---|---|
| 1a/1b | 21.8 | 18.8 |
| 2a | 5.2 | 9.8 |
| 2b | 7.6 | 11.0 |
| 3 | 8.7 | 4.7 |
| 4 | 8.9 | 8.8 |
| 5 | 47.9 | 46.9 |

% Activity compared to PBP5

| PBP | PY79 | DK694 |
|---|---|---|
| 1a/1b | 45.4 | 40.0 |
| 2a | 10.9 | 20.8 |
| 2b | 15.8 | 23.5 |
| 3 | 18.2 | 10.0 |
| 4 | 18.6 | 18.8 |
| 5 | 100.0 | 100.0 |

**Table S1.** Relative peptidase activity of different PBPs in *B. subtilis* PY79 (wild type) *vs* DK694 (Δ*pbpH*).(1) Activity of individual PBPs was compared to total activity as well as PBP5, in wild type and mutant strain. We assumed that in the absence of PBPH, a class B transpeptidase, the activity of PBP5, the only D,D-carboxypeptidase in *B. subtilis*, would be least affected.

| | Ampicillin | | Methicillin | |
|---|---|---|---|---|
| | PY79 | DK694 | PY79 | DK694 |
| PBP1a/1b | <0.01 | <0.01 | >1000 | >1000 |
| PBP2a | 0.0036 | 0.0061 | <0.01 | 0.028 |
| PBP2b | 0.0094 | 0.02 | <0.1 | 0.049 |
| PBP3 | 0.042 | 0.19 | 9 | 12.1 |
| PBP4 | 0.0037 | 0.0068 | 0.11 | 0.35 |
| PBP5 | 0.12 | 0.76 | 148.8 | 127.2 |

**Table S2.** $IC_{50}$ values (μg/mL) for individual PBPs from wild type (PY79) and Δ*pbpH* (DK694) strains of *Bacillus subtilis* are compared. Average of two individual experiments are provided. An $IC_{50}$ of >1,000 was assigned when GraphPad Prism reported an ambiguous number and significant inhibition was not seen at 1,000 μg/mL. In cases where GraphPad Prism resulted in ambiguous values, $IC_{50}$ was reported as lower than the concentration that caused >50% inhibition of that PBP.

**Table S3.** PBP inhibition profile shown by β-lactams at the MIC concentration.

| β-Lactam | MIC (μg/ml) | PBP(s) Completely Inhibited at MIC |
|---|---|---|
| **Aztreonam** | 16 | 1 |
| **Faropenem** | 0.125 | 1 |
| **Doripenem** | 0.0625 | none |
| **Meropenem** | 0.0625 | none |
| **Mecillinam** | 16 | 2a, 2b |
| **Penicillin V** | 32 | 2a, 2b, 3, 4, 5 |
| **Penicillin G** | 16 | 2a, 2b, 3, 4, 5 |
| **Ampicillin** | 8 | 2a, 2b, 3, 4 |
| **Amoxicillin** | 4 | 2a, 2b, 3, 4 |
| **Piperacillin** | 64 | 1, 2a, 2b, 3, 4 |
| **Methicillin** | 0.25 | 2a, 2b |
| **Oxacillin** | 0.25 | 4 |
| **Cloxacillin** | 0.125 | 4 |
| **6-APA** | 8-256 | none |
| **Cephalexin** | 0.5 | 2a, 2b |
| **Cefuroxime** | 0.125 | 1, 4 |
| **Cefotaxime** | 0.0625 | 1, 2b, 4 |
| **Ceftriaxone** | 0.25 | 1, 4 |
| **Cephalothin** | 0.03125 | 1, 4 |
| **Cefsulodin** | 8 | 1, 4 |
| **Cefoxitin** | 1 | 1, 2a, 2b, 4, 5 |

**Table S4.** Inhibition of Boc-FL labeling of PBPs in *B. subtilis* PY79.

| β-Lactam | Concentration[a] (μg/ml) | Relative % Boc-FL Labeling[b] PBP1a/1b | PBP2a | PBP2b/H | PBP3 | PBP4 | PBP5 |
|---|---|---|---|---|---|---|---|
| **Monobactam** | | | | | | | |
| **Aztreonam** | 0.01 | **21** | 104 | 99 | 106 | 107 | 104 |
| **Penem** | | | | | | | |
| **Faropenem** | 0.001 | **24** | 83 | 83 | 99 | 82 | 103 |
| **Carbapenem** | | | | | | | |
| **Doripenem** | 1 | 75 | **50** | **39** | **3** | 76 | 13 |
| **Meropenem** | 1 | 136 | 109 | 85 | **22** | 60 | **12** |
| **Penicillin** | | | | | | | |
| **Mecillinam** | 1 | 99 | 86 | **48** | 88 | 95 | 94 |
| **Penicillin V** | 0.001 | 64 | **4** | 62 | 105 | 56 | 101 |
| **Penicillin G** | 0.01 | 66 | **22** | 64 | 97 | **50** | 95 |
| **Ampicillin** | 0.001 | **24** | **1** | **4** | 93 | **7** | 92 |
| **Amoxicillin** | 0.1 | **34** | **30** | **42** | 77 | **32** | 91 |
| **Piperacillin** | 0.01 | **8** | 85 | 88 | 105 | 73 | 93 |
| **Methicillin** | 0.001 | 64 | **42** | 58 | 109 | 96 | 110 |
| **Oxacillin** | 0.01 | 60 | 70 | 68 | 87 | **48** | 95 |
| **Cloxacillin** | 0.1 | 67 | **47** | 64 | 100 | **10** | 78 |
| **6-APA** | 100 | 111 | 53 | 77 | 83 | 70 | **15** |
| **Cephalosporin** | | | | | | | |
| **Cephalexin** | 0.1 | 63 | **7** | **23** | 112 | 67 | 111 |
| **Cefuroxime** | 0.01 | **2** | 96 | 73 | 100 | 65 | 96 |
| **Cefotaxime** | 0.01 | **19** | 89 | 61 | 90 | **50** | 90 |
| **Ceftriaxone** | 0.01 | **15** | 88 | 85 | 85 | 76 | 90 |
| **Cephalothin** | 0.01 | **7** | **27** | **25** | 103 | **17** | 102 |
| **Cefsulodin** | 0.1 | **2** | 117 | 89 | 114 | 68 | 113 |
| **Cefoxitin** | 0.01 | **37** | 98 | 94 | 108 | **41** | 92 |

[a] Concentration of antibiotic required to reduce the subsequent labeling of Boc-FL to ≥50% for at least one PBP

[b] Inhibition of ≥50% of Boc-FL labeling is indicated by bolded text. Average of two independent experiments.

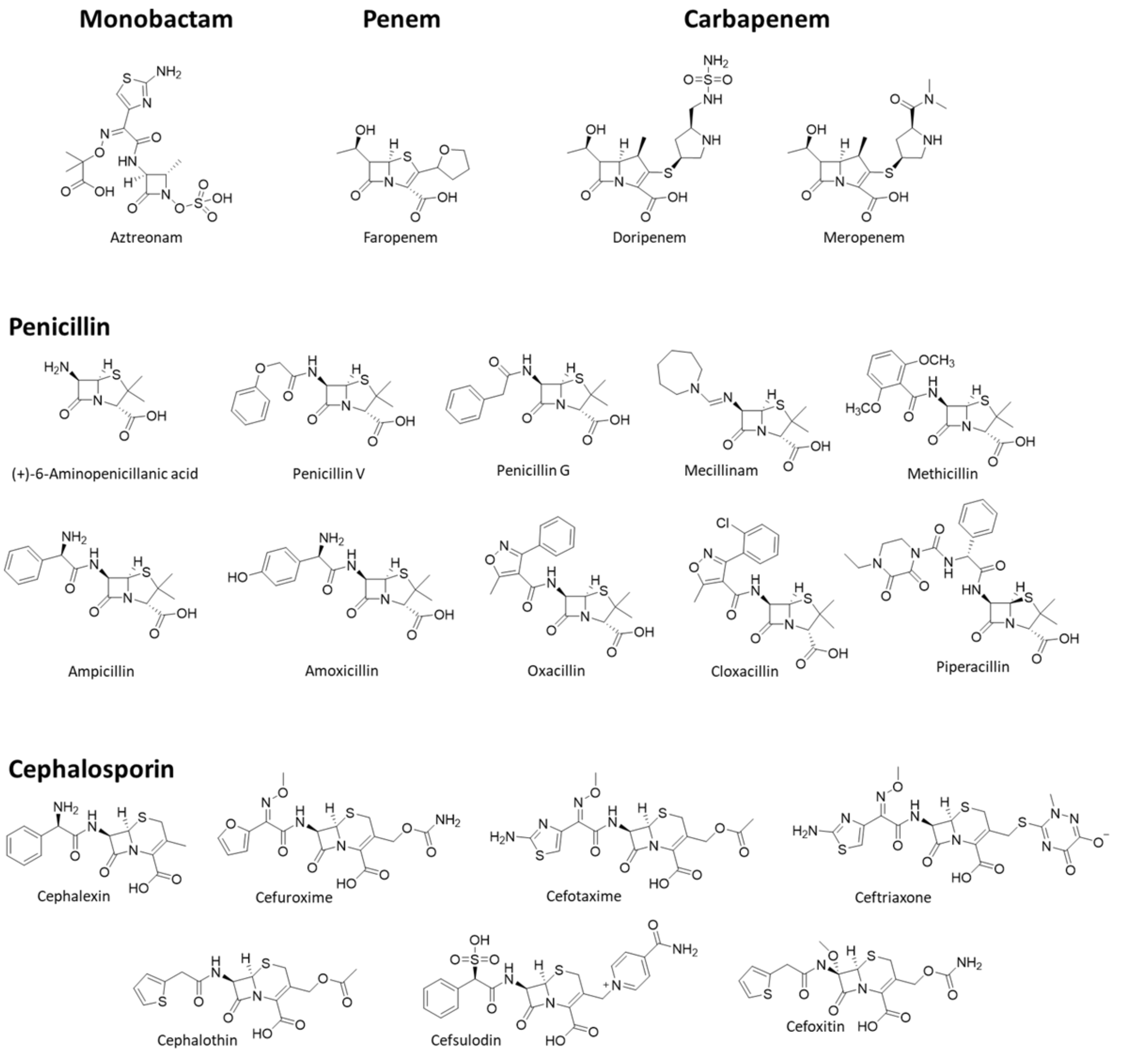


**Figure S1**. Structure of β-lactam antibiotics tested in this work.

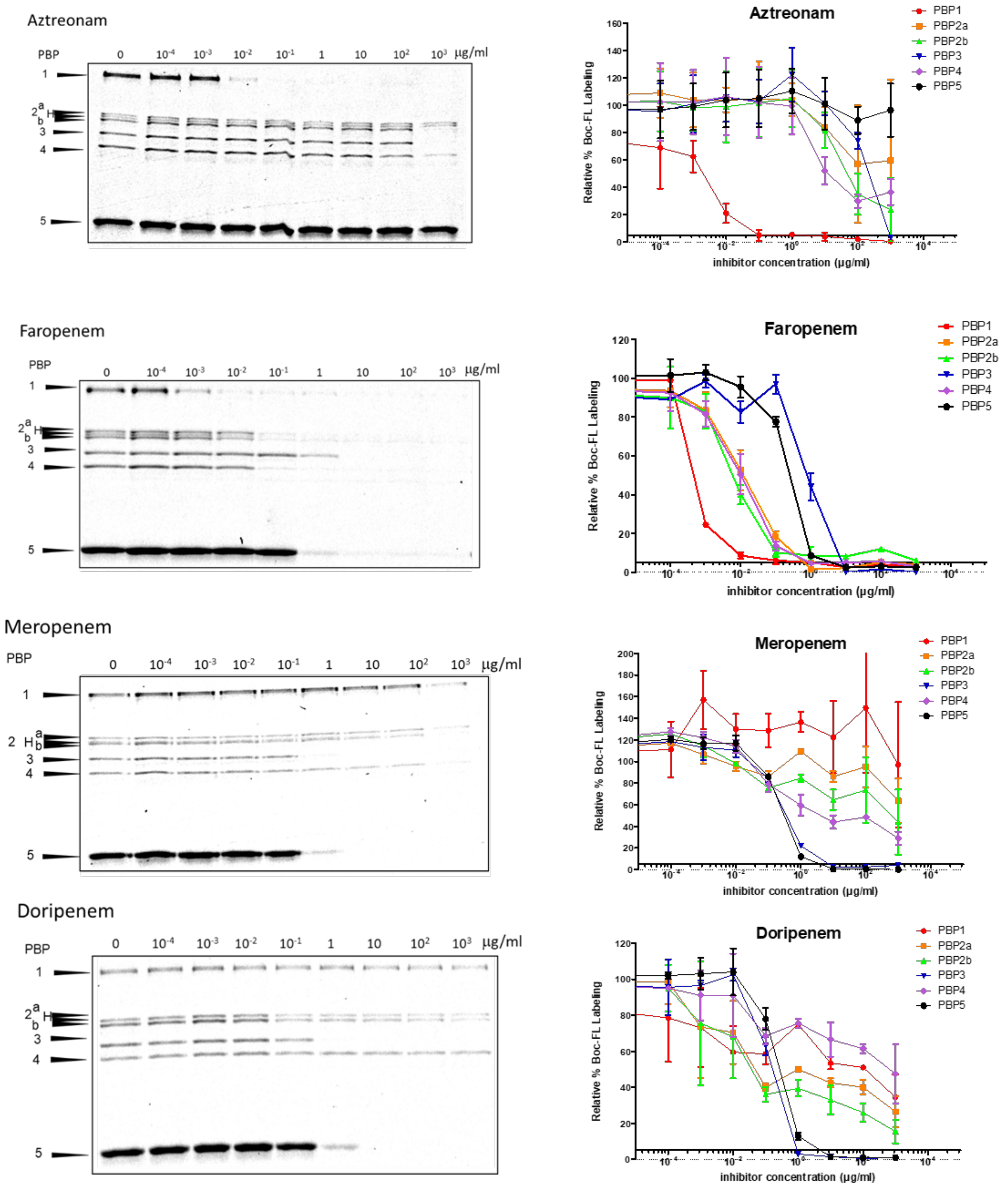
Aztreonam
PBP
0 10-4 10-3 10-2 10-1 1 10 10² 10³ μg/ml
1
2 a b
3
4
5
Aztreonam
PBP1
PBP2a
PBP2b
PBP3
PBP4
PBP5
Relative % Boc-FL Labeling
inhibitor concentration (μg/ml)
Faropenem
PBP
0 10-4 10-3 10-2 10-1 1 10 10² 10³ μg/ml
1
2 a b
3
4
5
Faropenem
PBP1
PBP2a
PBP2b
PBP3
PBP4
PBP5
Relative % Boc-FL Labeling
inhibitor concentration (μg/ml)
Meropenem
PBP
0 10-4 10-3 10-2 10-1 1 10 10² 10³ μg/ml
1
2 a b
3
4
5
Meropenem
PBP1
PBP2a
PBP2b
PBP3
PBP4
PBP5
Relative % Boc-FL Labeling
inhibitor concentration (μg/ml)
Doripenem
PBP
0 10-4 10-3 10-2 10-1 1 10 10² 10³ μg/ml
1
2 a b
3
4
5
Doripenem
PBP1
PBP2a
PBP2b
PBP3
PBP4
PBP5
Relative % Boc-FL Labeling
inhibitor concentration (μg/ml)

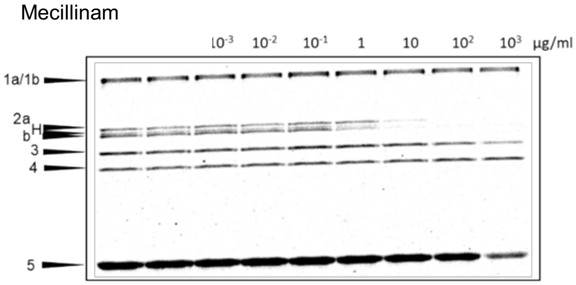
Mecillinam
$10^{-3}$ $10^{-2}$ $10^{-1}$ 1 10 $10^{2}$ $10^{3}$ µg/ml
1a/1b
2a
b
3
4
5

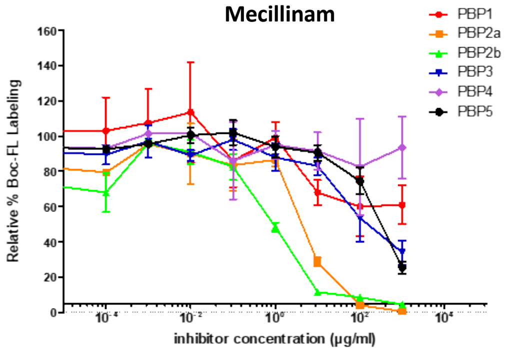
Mecillinam
PBP1
PBP2a
PBP2b
PBP3
PBP4
PBP5
Relative % Boc-FL Labeling
inhibitor concentration (µg/ml)

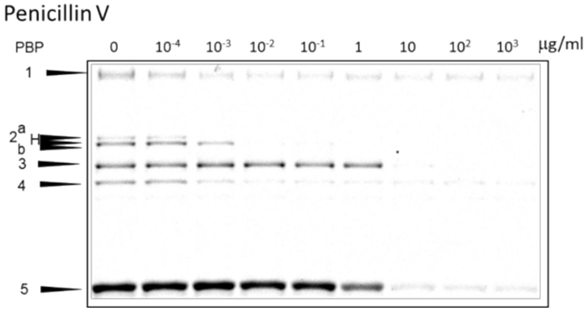
Penicillin V
PBP
0 $10^{-4}$ $10^{-3}$ $10^{-2}$ $10^{-1}$ 1 10 $10^{2}$ $10^{3}$ µg/ml
1
2a
b
3
4
5

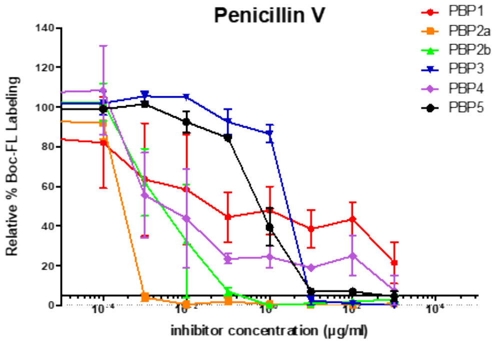
Penicillin V
PBP1
PBP2a
PBP2b
PBP3
PBP4
PBP5
Relative % Boc-FL Labeling
inhibitor concentration (µg/ml)

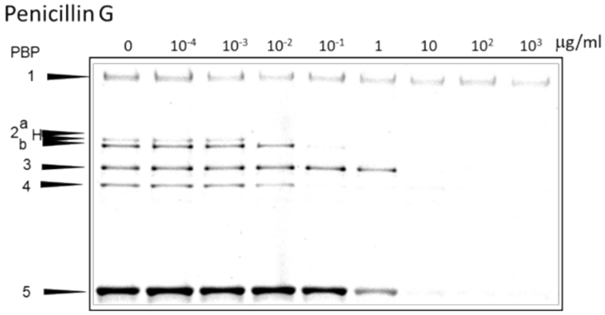
Penicillin G
PBP
0 $10^{-4}$ $10^{-3}$ $10^{-2}$ $10^{-1}$ 1 10 $10^{2}$ $10^{3}$ µg/ml
1
2a
b
3
4
5

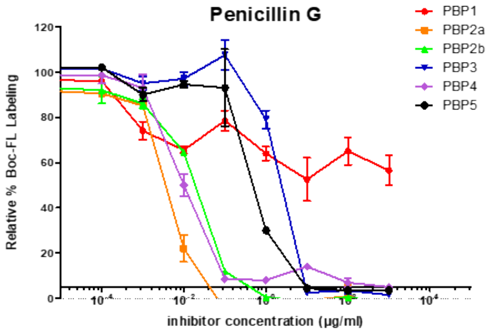
Penicillin G
PBP1
PBP2a
PBP2b
PBP3
PBP4
PBP5
Relative % Boc-FL Labeling
inhibitor concentration (µg/ml)

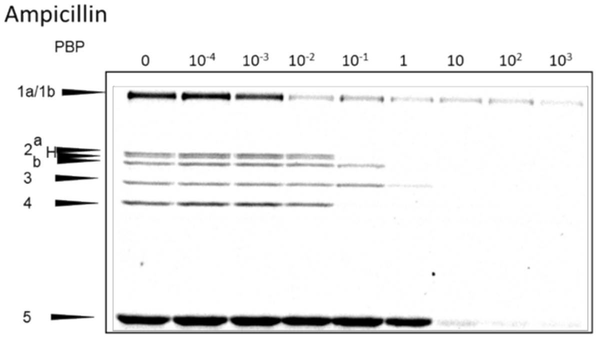
Ampicillin
PBP
0 $10^{-4}$ $10^{-3}$ $10^{-2}$ $10^{-1}$ 1 10 $10^{2}$ $10^{3}$
1a/1b
2a
b
3
4
5

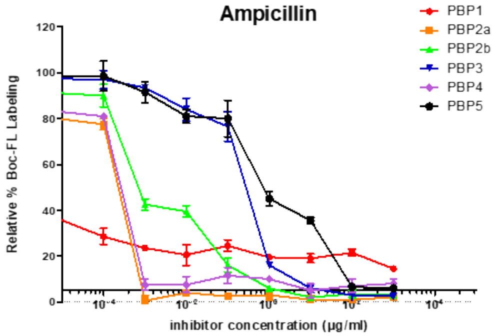
Ampicillin
PBP1
PBP2a
PBP2b
PBP3
PBP4
PBP5
Relative % Boc-FL Labeling
inhibitor concentration (µg/ml)

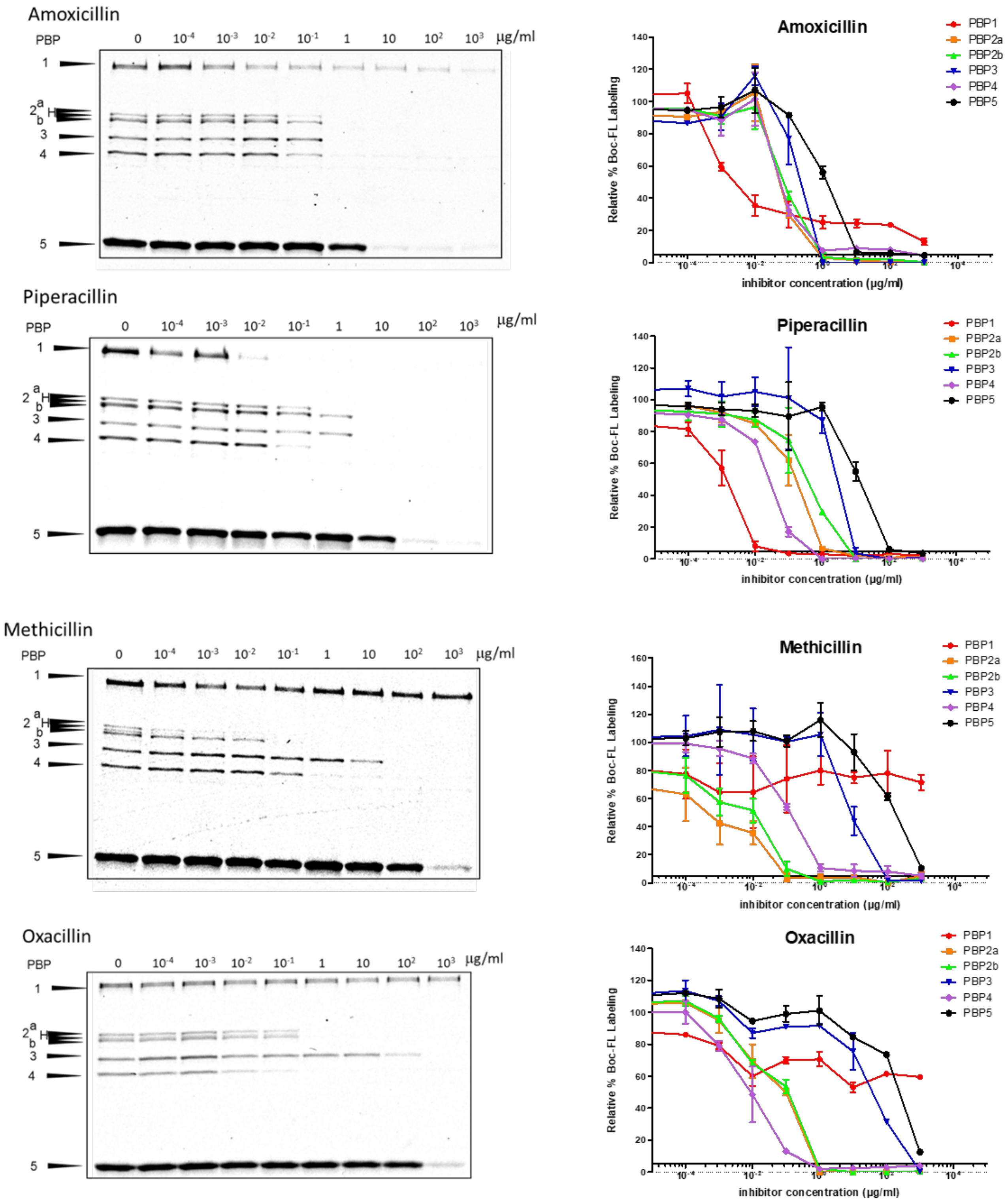

Amoxicillin
PBP
0
10-4
10-3
10-2
10-1
1
10
102
103
μg/ml
1
2 a b
3
4
5
Amoxicillin
Relative % Boc-FL Labeling
inhibitor concentration (μg/ml)
PBP1
PBP2a
PBP2b
PBP3
PBP4
PBP5
Piperacillin
Piperacillin
Methicillin
Methicillin
Oxacillin
Oxacillin

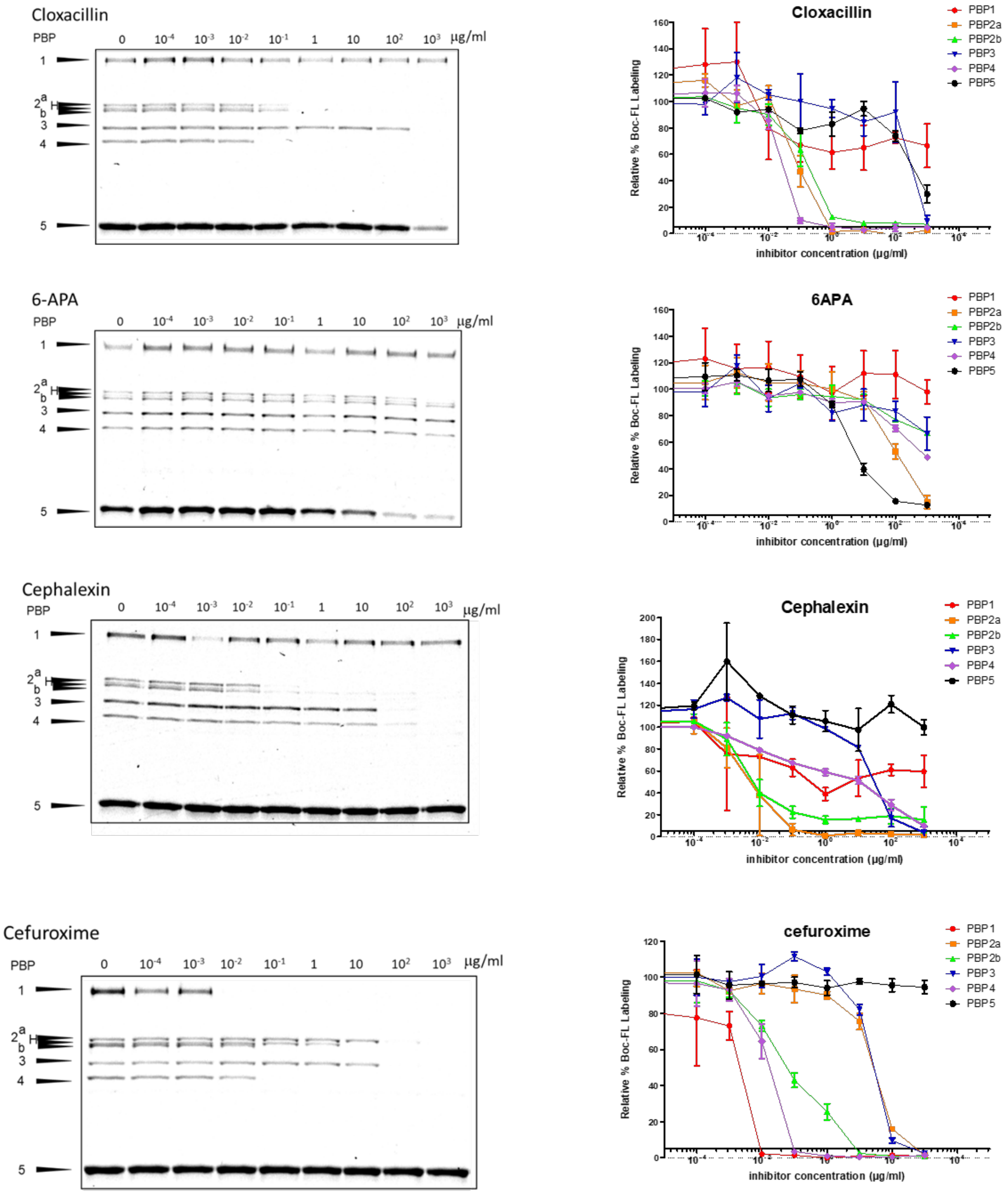
Cloxacillin
PBP
0
10-4
10-3
10-2
10-1
1
10
102
103
µg/ml
1
2
a
b
3
4
5
Cloxacillin
PBP1
PBP2a
PBP2b
PBP3
PBP4
PBP5
Relative % Boc-FL Labeling
inhibitor concentration (µg/ml)
6-APA
6APA
Cephalexin
Cefuroxime
cefuroxime

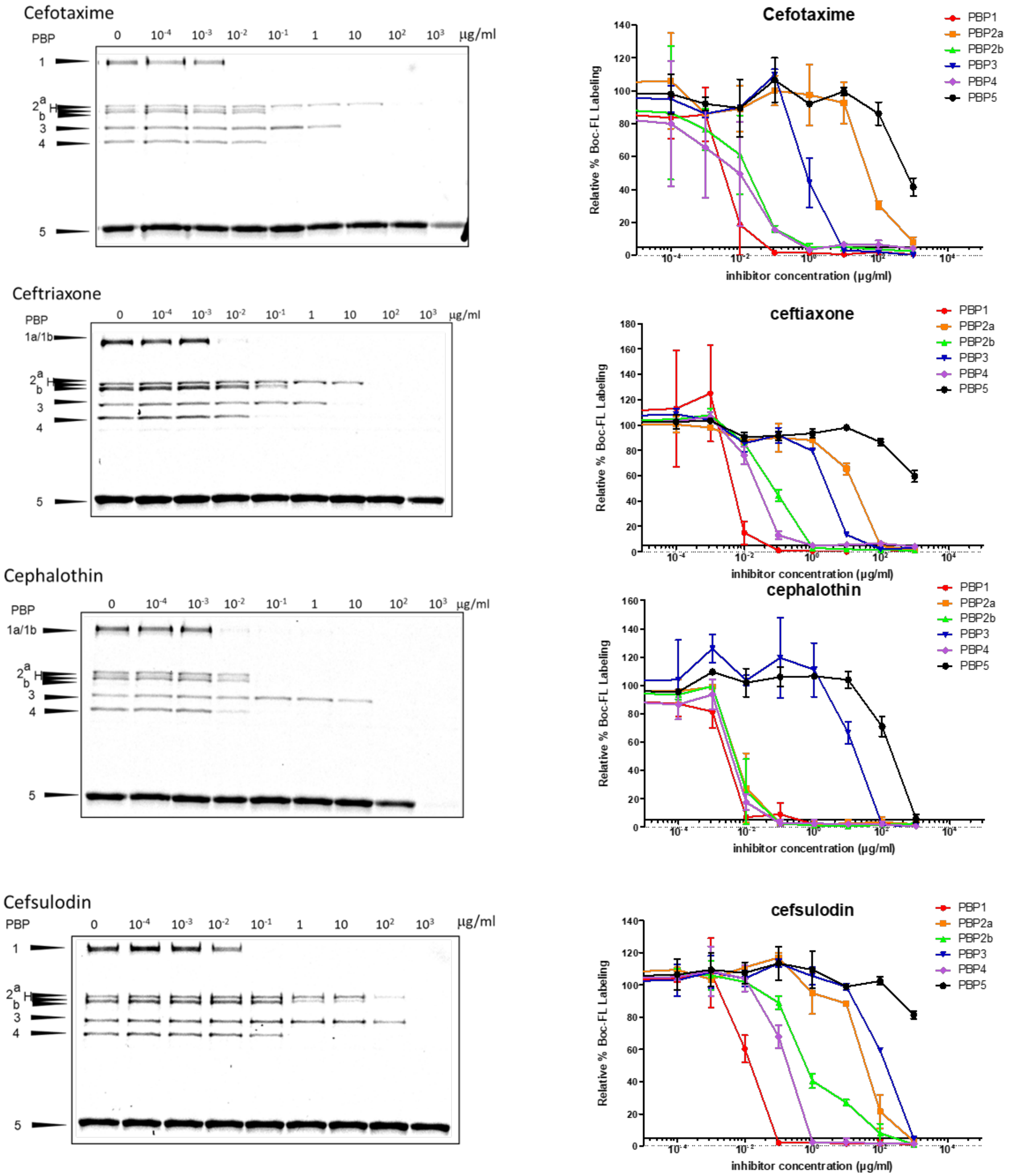
Cefotaxime
PBP
0 10-4 10-3 10-2 10-1 1 10 102 103 μg/ml
1
2 a b
3
4
5
Cefotaxime
PBP1
PBP2a
PBP2b
PBP3
PBP4
PBP5
Relative % Boc-FL Labeling
inhibitor concentration (μg/ml)
Ceftriaxone
PBP
1a/1b
ceftiaxone
Cephalothin
cephalothin
Cefsulodin
cefsulodin

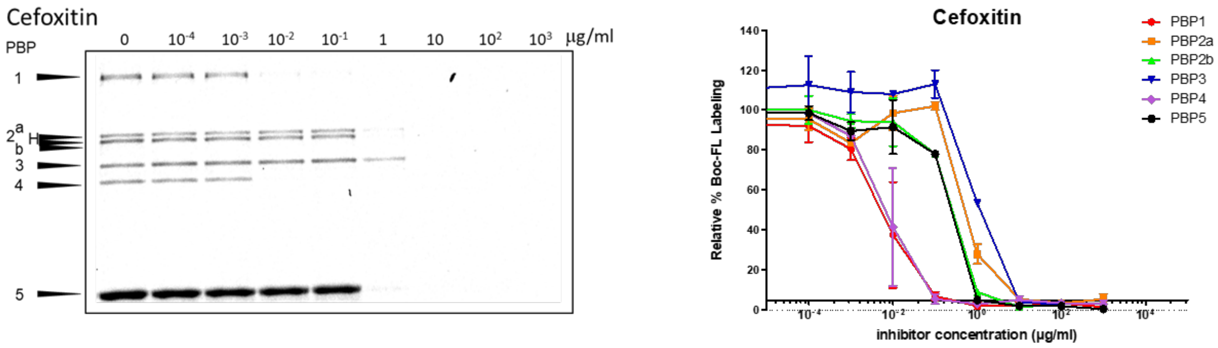


**Figure S2.** Representative SDS-PAGE gels (one of two independent experiments) and the representative graphs for β-lactam titration of the PBPs in *B. subtilis* PY79. Whole cells were treated with various concentrations of antibiotics and subsequently labeled with 5 μg/ml BOCILLIN FL (Boc-FL). Gel bands were integrated and relative fluorescence intensity was calculated in comparison with the control. The average relative intensity from two independent assays is plotted against β-lactam concentration.

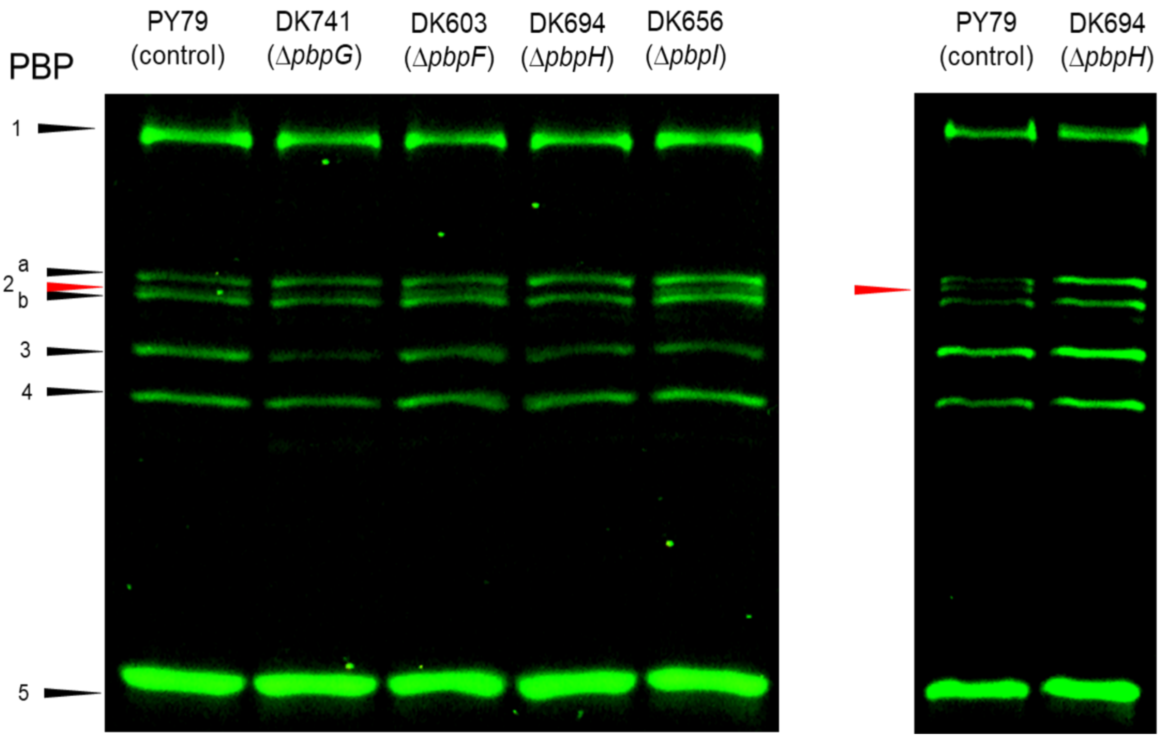


**Figure S3**. Left: PBP activity profile of *B. subtilis* PY79 (wild type) and mutant strains.(1) Cells were treated with 5 μg/mL Bocillin-FL for 10 minutes at RT. *pbpG*, *pbpF*, *pbpH* and *pbpI* encode PBPs 2d, 2c, H and 4b, respectively. All strains, except Δ*pbpH*, display an additional faint band between PBP2a and 2b (pointed out by a red arrow). Lysate protein concentration was adjusted at 5.0 mg/mL.

Right: PBP activity profile of *B. subtilis* PY79 (wild type) and DK694 (Δ*pbpH*) compared. Lysate protein concentration was adjusted at 3.0 mg/mL to better resolve bands in PBP2 region.

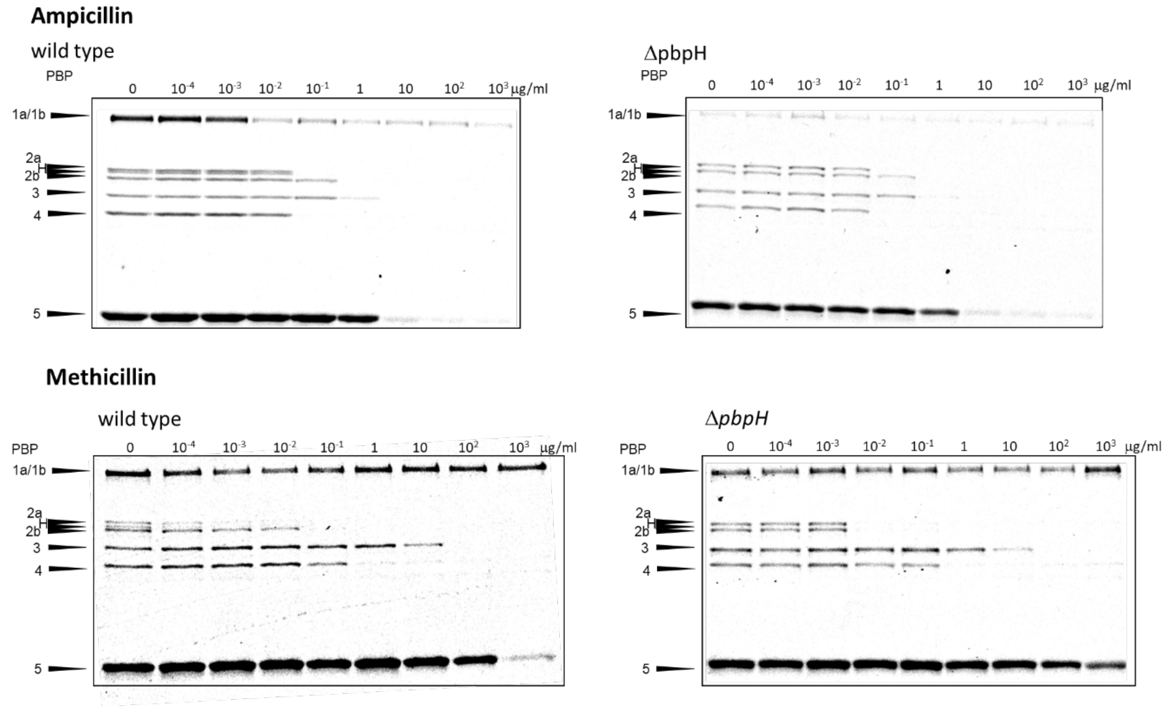


**Figure S4**. Representative SDS-PAGE gels (one of two independent experiments) for ampicillin (top) and methicillin (bottom) titration of PBPs in *B. subtilis* DK694 (Δ*pbpH*) compared with wild-type. Whole cells were treated with various concentrations of antibiotics and subsequently labeled with 5 µg/mL BOCILLIN FL (Boc-FL).

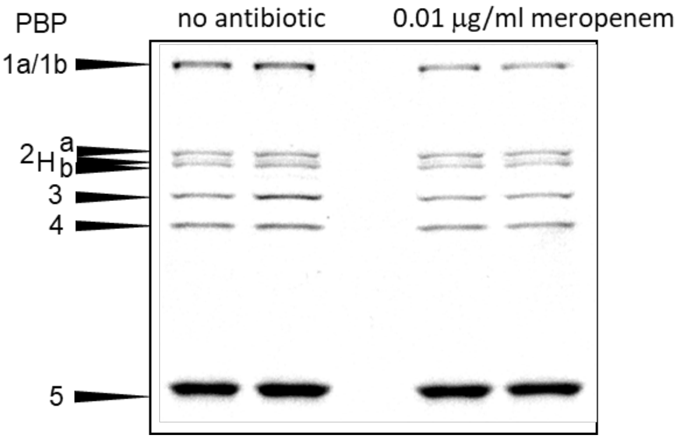


**Figure S5**. PBP activity profile of *B. subtilis PY79* cells grown in the presence of 0.01 µg/mL meropenem after 2 hours. Cells from antibiotic-treated culture and control were harvested at the same time. Boc-FL labeling was performed using the described method.

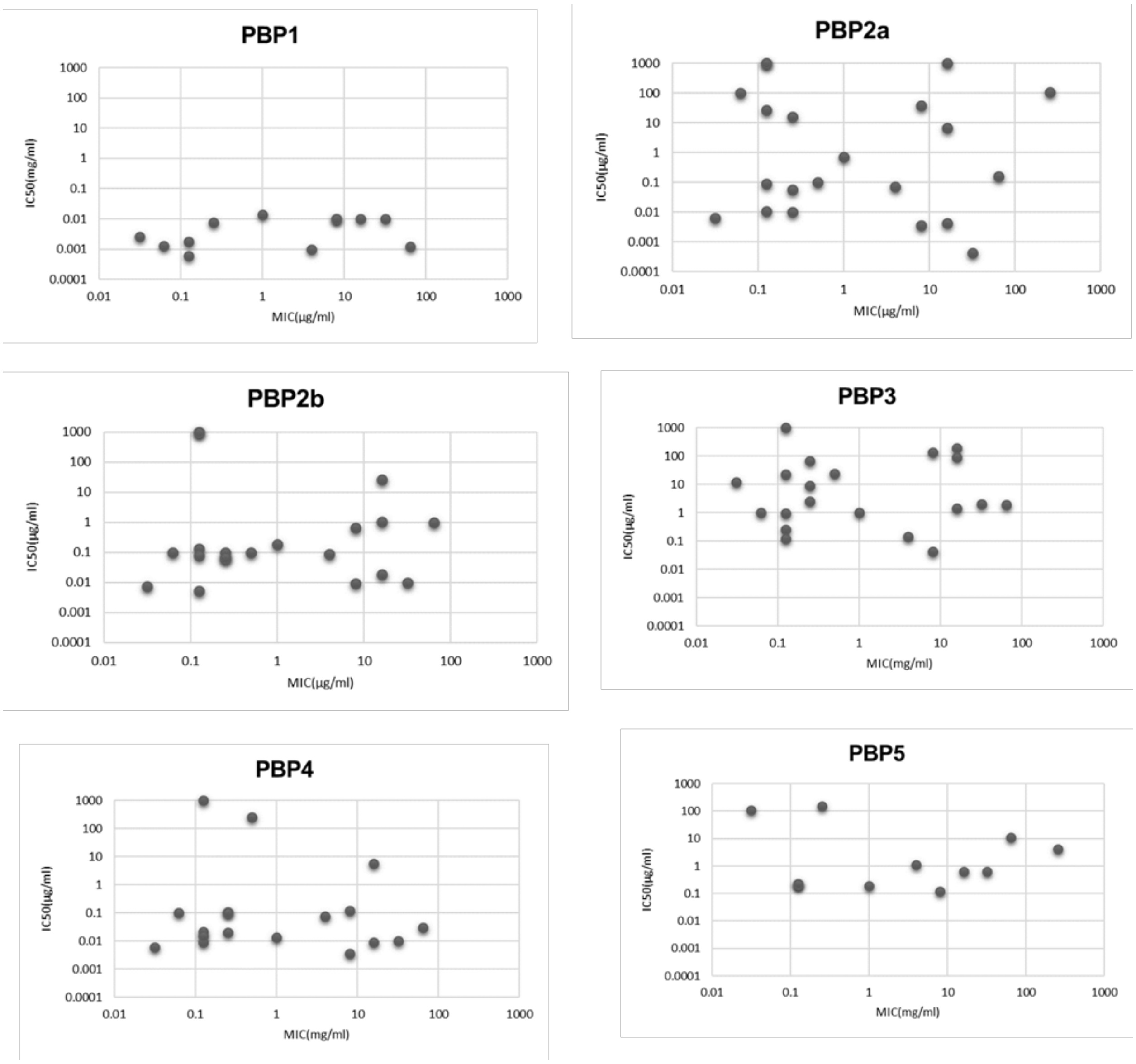


**Figure S6.** Graphs of MIC *vs* $IC_{50}$ for PBPs in *B. subtilis*.

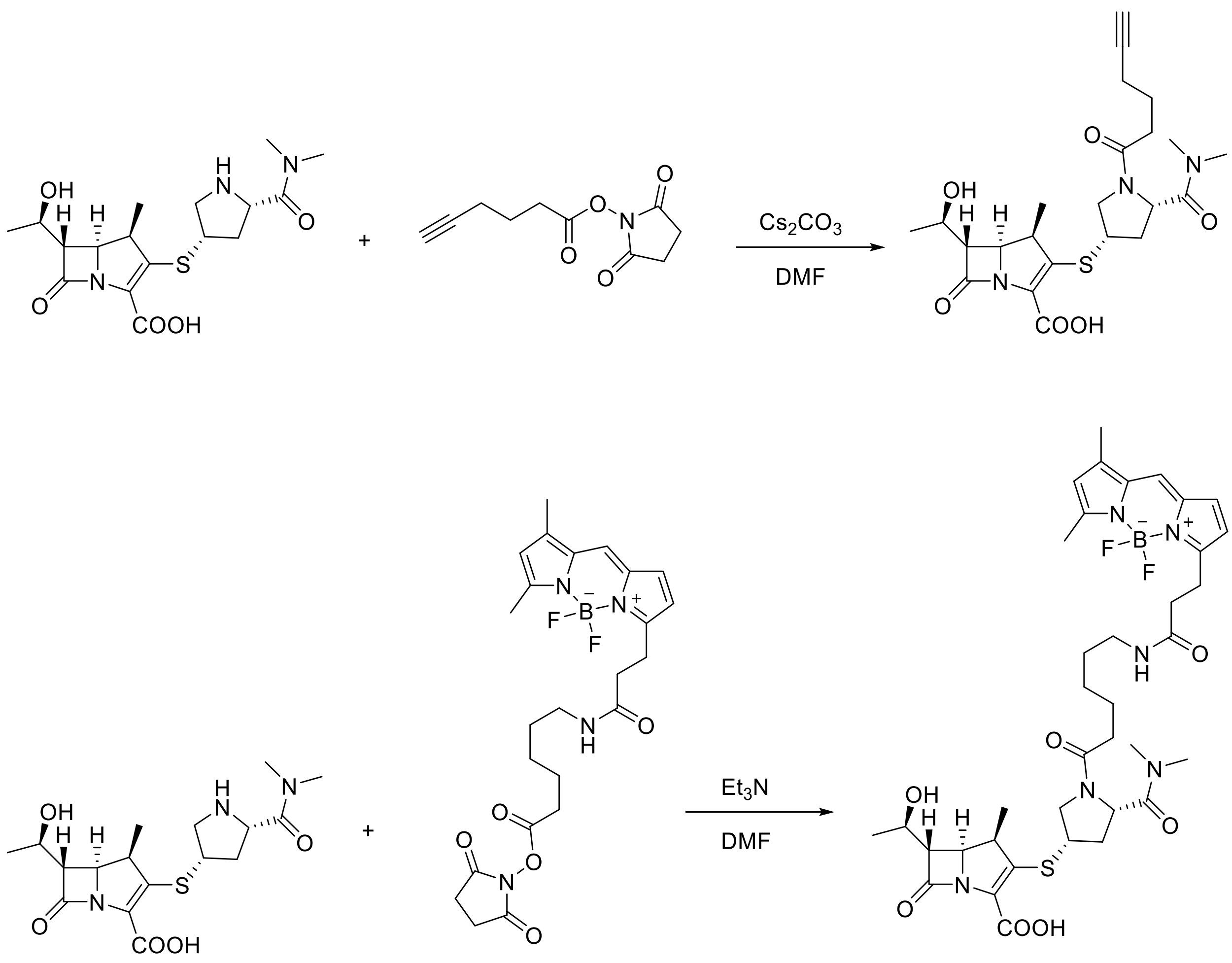


**Scheme S1.** General synthetic scheme for meropenem-based probes

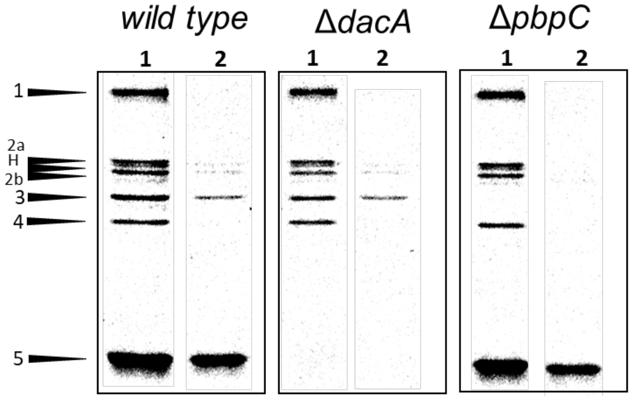


**Figure S7.** PBP labeling profile of MEM-BODIPY in *B. subtilis* 3610 (wild type), DK654 (Δ*dacA*) and DK695 (Δ*pbpC*). **1** : Boc-FL 10μM; **2** : mem-BODIPY 2μM.

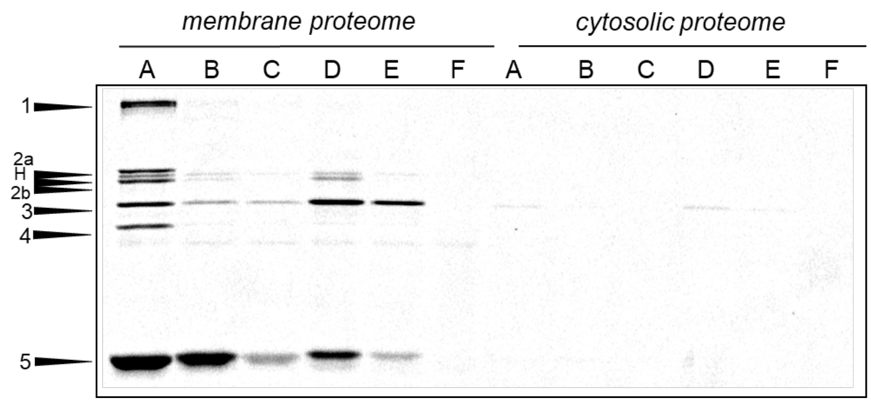


**Figure S8.** Meropenem targets in membrane and cytosolic fractions of *B. subtilis 3610* strain.

**A** : Boc-FL 10uM

**B** : Mem-BODIPY 0.5uM

**C** : Mem-BODIPY 0.5uM + 6APA 100ug/mL

**D** : Click-mem 5uM → AF488-Az

**E** : Click-mem 5uM + 6APA 100ug/mL→ AF488-Az

**F** : PBS→ AF488-Az

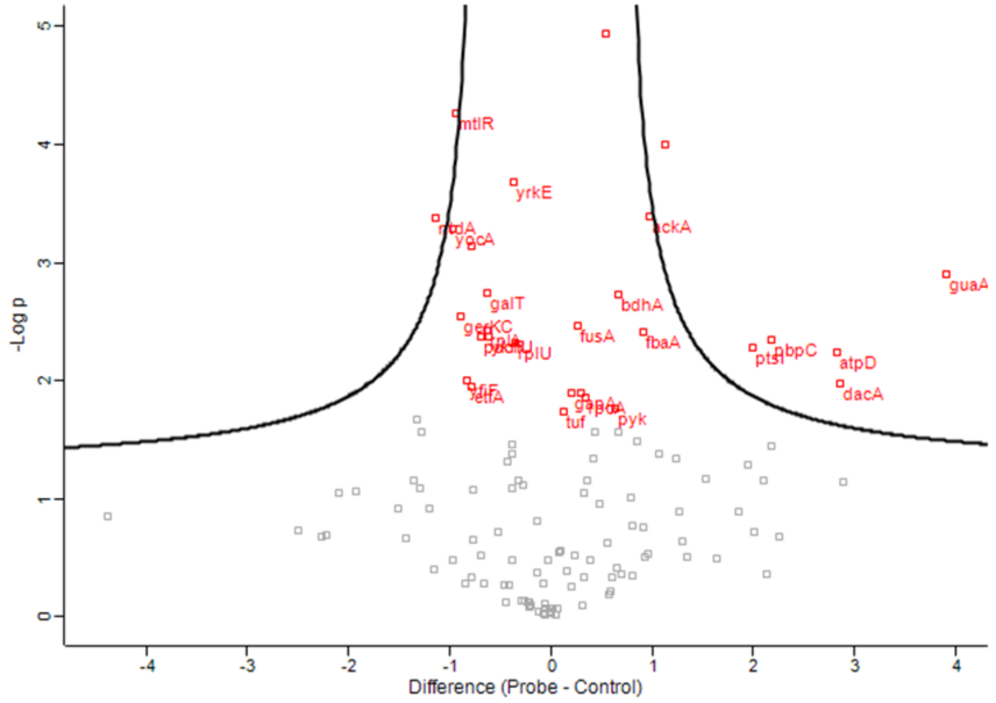


**Figure S9.** Volcano plot of proteins enriched by click-MEM probe. PBP3 and PBP5, encoded by *pbpC* and *dacA* genes respectively, are among the detected proteins. Following parameters were used for T-test: FDR = 0.01, S0 = 2.0; Volcano plot: FDR = 0.05, S0 = 0.3.

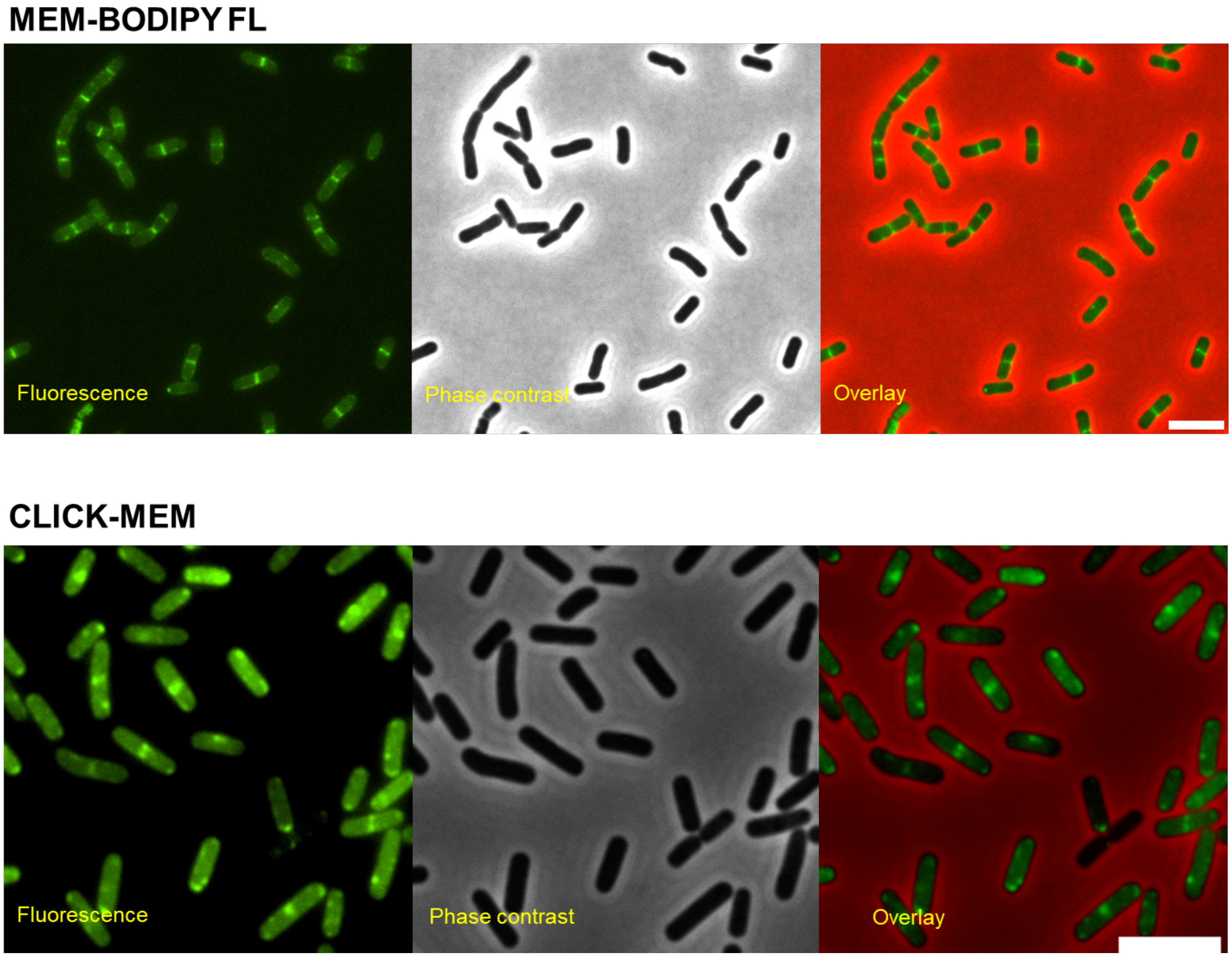


**Figure S10.** Visualization of PBP3 activity in live *B. subtilis DK654* cells. Exponentially growing *B. subtilis DK654* cells were labeled with 2 µM MEM-BODIPY FL (*top*) or 5 µM Click-MEM followed by click reaction with 50 µM AlexaFluor488-azide in the presence of 2.5 mM sodium ascorbate, 200 µM BTTAA and 100 µM copper sulfate in minimal media (*bottom*). Further details provided in the experimental protocol. Scale bar = 5 µm.

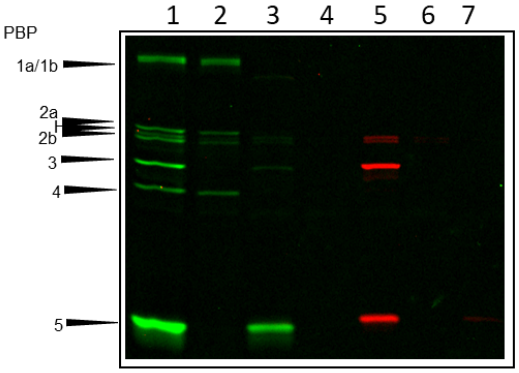


1 : PBS → Boc-FL 10 µM
2 : 10 µg/ml meropenem → Boc-FL 10 µM
3 : PBS → MEM-BODIPY FL 2 µM
4 : 10 µg/ml meropenem → MEM-BODIPY FL 2 µM
5 : PBS → click-MEM 5 µM → TAMRA-azide
6 :10 µg/ml meropenem → click-MEM 5 µM → TAMRA-azide
7 : PBS→ TAMRA-azide

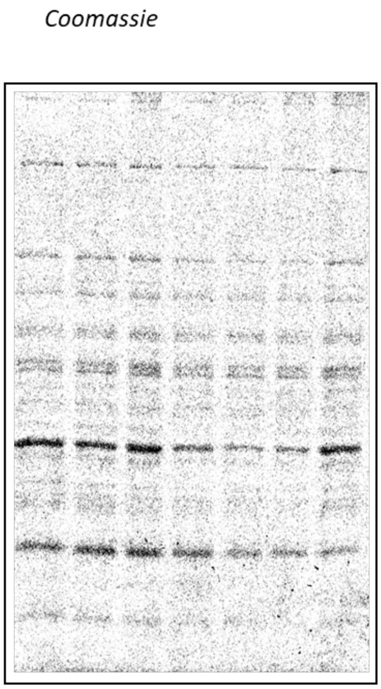


**Figure S11.** Meropenem pretreatment inhibits PBP3 and PBP5 labeling by MEM-probes. Cells were treated with 0 or 10 µg/ml meropenem for 30 min, washed and then labeled with 10 µM Boc-FL for 10 mins, or 2 µM MEM-BODIPY FL or 5 µM click-MEM for 30 min at R.T. Cells were lysed, membrane fraction collected and click reaction with TAMRA-azide was performed for click-MEM tagged samples for 40 mins at R.T.

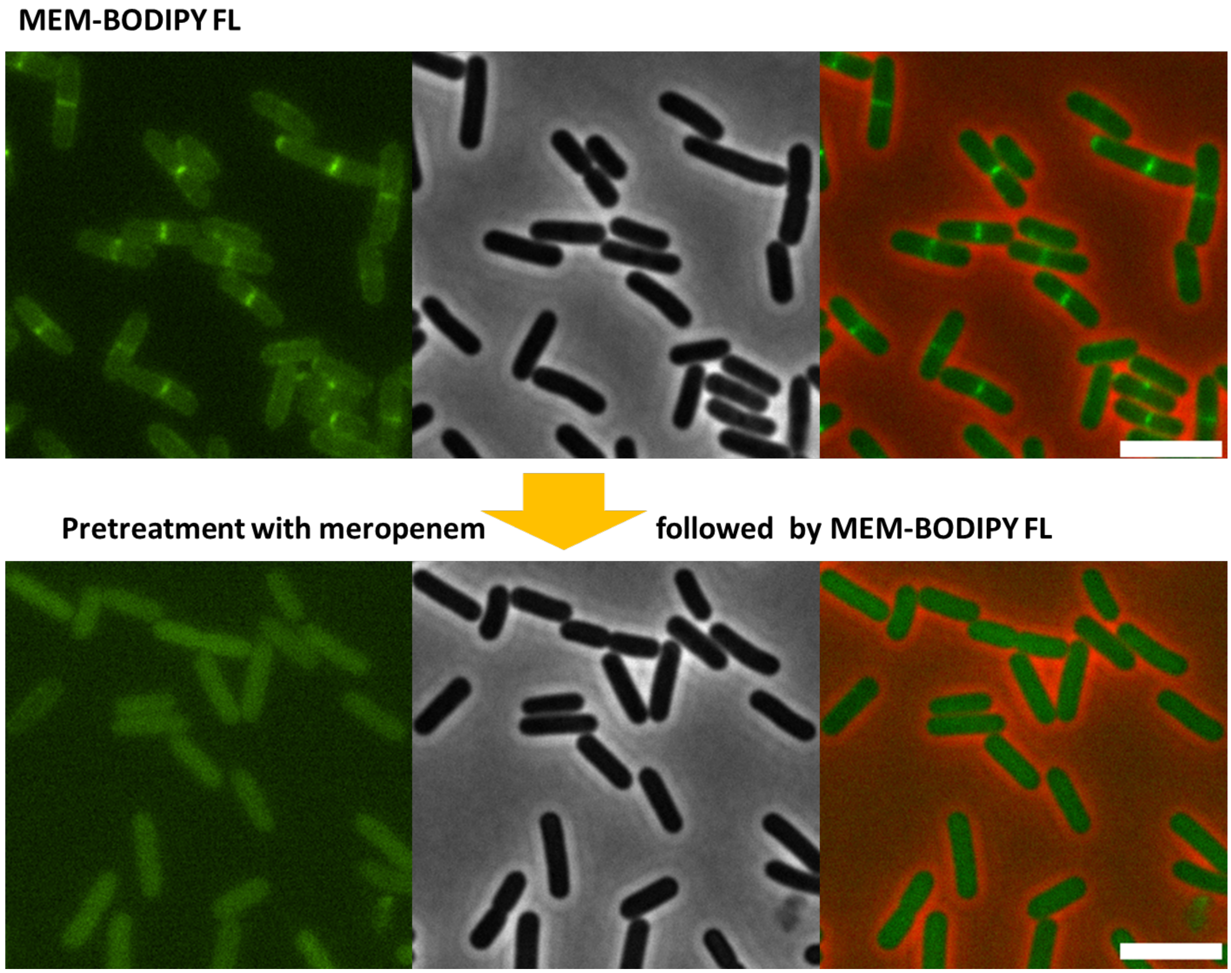


**Figure S12.** Preteatment with meropenem prevents PBP3 labeling by MEM-BODIPY FL probe. Exponentially growing *B. subtilis DK654* cells were treated with 10 µg/ml meropenem and incubated at R.T. for 30 min. Next, cells were labeled with 2 µM MEM-BODIPY FL and imaged at 100X magnification. Scale bar = 5 µm.

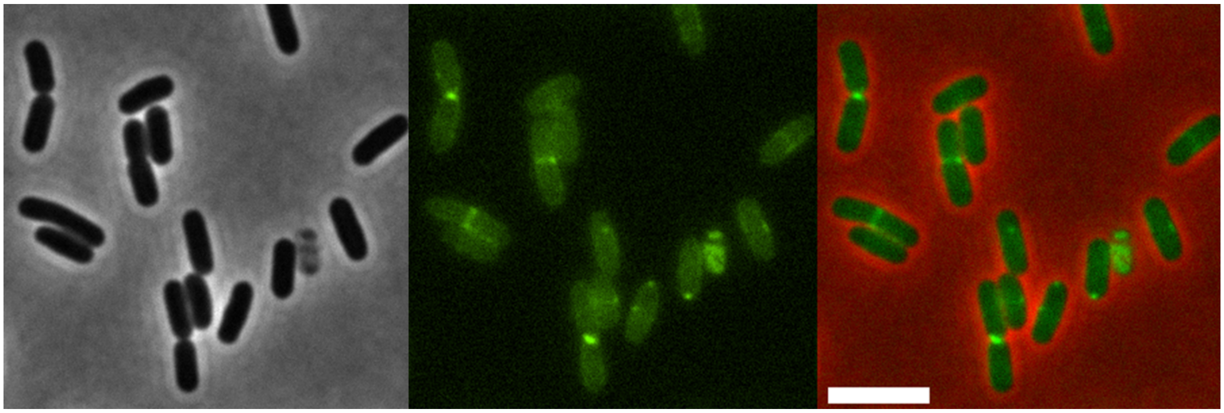

**Figure S13.** Visualization of PBP3 activity in live *B. subtilis DK654* cells pretreated with mecillinam. Exponentially growing *B. subtilis DK654* cells were treated with 5 µg/ml mecillinam and incubated at R.T. for 30 min. Next, cells were labeled with 2 µM MEM-BODIPY FL and imaged at 100X magnification. Scale bar = 5 µm.

**General Materials and Methods**

β-lactam compounds were selected from different subclasses including penicillins, cephalosporins, carbapenems, penems and monobactams. Faropenem, doripenem, meropenem, (+)-6-aminopenicillanic acid (6-APA), ampicillin, methicillin, oxacillin, cloxacillin, piperacillin, cephalexin, cefsulodin, cefoxitin, cephalothin, cefuroxime, ceftriaxone and cefotaxime were purchased from Sigma-Aldrich (St. Louis, MO). Penicillin V and penicillin G were from USP. Mecillinam was purchased from Fluka. Aztreonam and amoxicillin were obtained from MP Biomedicals (Solon, OH). Bocillin-FL was purchased from Life Technologies (Grand Island, NY).

All antibiotics were stored as solids at 4 °C (except for meropenem, doripenem, aztreonam and faropenem which were stored at -20 °C) and were dissolved in Milli-Q purified water (unless noted otherwise) at 10 mg/ml immediately before each experiment. 6-APA, amoxicillin, and cefuroxime were not soluble in water. Thus, they were dissolved in phosphate-buffered saline (PBS; 8.00 g/L NaCl, 0.20 g/L KCl, 1.44 g/L $Na_2HPO4$, 0.24 g/L $KH_2PO4$; pH 7.4) at a 1 mg/mL concentration. All stock solutions were serially diluted (10-fold) in PBS to make 0.0001 to 1,000 mg/mL working solutions. For MIC determination studies, antibiotics were dissolved and serially diluted in LB broth.

All reactions were carried out in anhydrous solvents under Ar atmosphere. Hydroxybenzotriazole (HOBt) was purchased from Chem-Impex International, *N*-(3-Dimethylaminopropyl)-N-ethylcarbodiimide HCl (EDC•HCl) and ammonium formate were purchased from Fluka. BODIPY-succinimidyl ester was purchased from ThermoFischer Company. Meropenem (anhydrous) was purchased from ChemCruz. Oxalyl chloride was purchased from Acros. All other reagents were purchased from Sigma-Aldrich and used without further purification.

AFDye 488 azide (equivalent of Alexa Fluor ® 488 azide), biotin-PEG3-azide, and (4-{[bis-(1-tert-butyl-1H-[1,2,3]triazol-4-ylmethyl)-amino]-methyl}-[1,2,3]triazol-1-yl)-acetic acid (BTTAA) were purchased from Click Chemistry tools. Tris(2-carboxyethyl)phosphine (TCEP), Tris[(1-benzyl-1H-1,2,3-triazol-4-yl)methyl]amine (TBTA), sodium ascorbate and copper sulfate were purchased from Sigma-Aldrich. All click reagents were kept at -20 °C as powder, except TCEP (stored at 4 °C) and copper sulfate and sodium ascorbate which were stored at room temperature. Stock solutions (5 mM) of azide cargo were prepared in DMSO and stored at -20 °C. All other solutions were prepared fresh by dissolving the required amounts in MilliQ water.

$^{1}H$ and $^{13}C$ NMR spectra were recorded on HD-500 spectrometers and are reported in parts per million (δ). The following abbreviations were used to explain the multiplicities: s = singlet, d = doublet, t = triplet, q = quartet, quin = quintet, m = multiplet. UPLC-ESI-TOF MS instrumentation (Agilent, 6220) was used to obtain accurate mass spectral data. HPLC purification was performed using an Agilent 1200 HPLC using a reverse phase column (Agilent Eclipse plus C18, 5 µm, 150 × 4.6 mm) detected by diode array detector (200–600 nm). Gradients consisted of 5–95% B (A: $H_2O$, 0.1% formic acid (FA); B: $CH_3CN$, 0.1% FA) over 20-30 minutes.

**Bacterial Culture and PBP Labeling**

**Culture growth.** *B. subtilis* PY79 is a prototrophic laboratory strain and has been commonly exploited for studying different cellular pathways.(1, 2) *B. subtilis* PY79 and all the other strains(3) were cultured in Luria-Bertani (LB) broth at 37 °C with shaking at 220 rpm to reach the desired optical density at 600 nm ($OD_{600}$).

**β-Lactam titration.** *B. subtilis* cells from 1.0 mL of an exponential culture at an $OD_{600}$ of 0.4 were harvested by centrifugation (16,000 × *g* for 2 min at room temperature). The cell pellets were washed in 1 mL of PBS (pH 7.4) by gently pipetting up and down a few times, and the cells were again pelleted. The same centrifugation settings were used throughout the assay, unless otherwise indicated. The cells were resuspended in 50 μL PBS containing 0.0001 to 1,000 μg/mL β-lactam antibiotics, and a reference sample was resuspended in 50 μL PBS without antibiotic (blank). After 30 min of incubation at room temperature, the cells were pelleted, washed in 1 mL PBS, and resuspended in 50 μL PBS containing 5 μg/mL Boc-FL (0.1% DMSO). After 10 min of incubation at room temperature, the cells were pelleted and washed in 1 mL PBS.

**MEM-BODIPY Probe Concentration Determination.** All probes were stored as DMSO solutions at -80 °C. Concentration of MEM-BODIPY was determined by measuring UV-Vis absorption of its solution at $\lambda_{max}$ of its corresponding fluorophore. 10X and 100X dilutions of probe was made in methanol and absorbance was read at 504 nm using NanoPhotometer P330 (IMPLEN). The average of 3 reads of the concentration that was in 0.1-0.9 range was used to calculate concentration. ($\varepsilon$ = 85,000 $M^{-1}.cm^{-1}$).

**Probe Labeling.** *B. subtilis* cells from 1.0 mL of an exponential culture at an $OD_{600}$ of 0.4 were harvested by centrifugation (16,000 × *g* for 2 min at room temperature). The cell pellets were washed in 1 mL of PBS (pH 7.4) by pipetting up and down a few times, and the cells were again pelleted using the same setting. Cells were subsequently suspended in 50 μL PBS containing 0.5-2 uM MEM-BODIPY or 1-10 uM click-MEM. After 30 min of incubation at room temperature, cells were pelleted, washed with 1 mL PBS, and pelleted again.

**Click Reaction.** *B. subtilis* cell pellets labeled by click-MEM (as described above) were suspended in 50 μL of click reaction cocktail consisting of PBS containing 20-50 μM AF488 azide, 1 mM TCEP, 0.5 mM TBTA and 1 mM $CuSO_4$. The resulting suspension was incubated at room temperature for an hour. For SDS PAGE analysis, cells were subsequently pelleted and washed with 1 mL PBS. Membrane proteome collection and analysis was performed as described for β-lactam titration.

For fluorescent imaging experiments, cell pellets were suspended in 50 μL of the click reaction cocktail consisting of S750 minimal media containing 50 μM AF488 azide, 2.5 mM sodium ascorbate, 0.2 mM BTTAA and 0.1 mM $CuSO_4$ and the incubation time was dropped to 30 min.

**Gel-Based Analysis.** Labeled cells were resuspended in 100 μL PBS containing 10 mg/ml lysozyme (chicken egg white, Fluka) and statically incubated at 37 °C for 30 min, after which samples were sonicated using a Hielscher vial tweeter UP200St (70% C, 95% A, 5% adjustment snap and 1s SD Interval / 10s for 6 x 1 min intervals with 1 min cooling time in between) on ice. The membrane proteome was isolated by centrifugation at 21,000 × *g* for 15 min at 4 °C, as previously reported(4, 5). The supernatant was discarded, the membrane was resuspended in 100 μL PBS and protein concentration adjusted to 5.0 mg/mL by dilution with PBS. The protein concentration was measured by NanoPhotometer P330 (IMPLEN). Thirty microliters of proteome sample was dispensed into a clean 1.5-mL microcentrifuge tube and 10 μL of 4 × SDS-PAGE loading buffer was added to each sample. The samples were heated for 5 min at 90-95 °C to denature the proteins then cooled to RT. Twelve microliters of sample were loaded onto a 10% SDS-PAGE gel (acrylamide:bis-acrylamide = 29:1). The protein bands were separated by gel electrophoresis for 1.5 h at 180 V, 400 mA, 60 W. The gel was rinsed with distilled water three times before fluorescence scanning using a Typhoon 9210 gel scanner (Amersham Biosciences,

Pittsburgh, PA) with a 526-nm wavelength short-pass filter at 50-μm resolution. The gel images were analyzed using ImageJ software (National Institutes of Health, Bethesda, MD). The background signal of the gel images was subtracted, and the brightness and contrast were adjusted to optimize the signal-to-noise ratio (all operations were uniformly performed over the entire gel). Integrated density values were measured for gel band quantitation, and Boc-FL labeling of each PBP in antibiotic-treated samples was shown relative to the blank. Integrated density values were inserted into GraphPad Prism (GraphPad Software, La Jolla, CA) to create graphs showing relative percentiles of Boc-FL labeling versus inhibitory concentrations (ICs). Determinations of 50% IC values ($IC_{50}$s) were performed using the same software. For all dose-response curves, data were fitted to a four-parameter logistic equation. The average of $IC_{50}$ values calculated from two independent assays was reported for individual PBPs for each β-lactam.

**Proteomic Analysis of Meropenem Targets**

**Protein Enrichment and Digestion.** *B. subtilis* membrane lysates labeled by click-MEM (as described above) were suspended in 50 μL of click reaction cocktail consisting of PBS containing 50 μM biotin-PEG3-azide (or Alexa488-Fluor azide as control for click reaction/binding), 1 mM TCEP, 0.5 mM TBTA and 1 mM $CuSO_4$. The resulting suspension was incubated at room temperature for one hour (fluorescent samples were run on 10% SDS PAGE gels to detect the tagged PBPs as described above). To remove excess biotin and click reagents, protein precipitation was performed using a ProteoExtract ® protein extraction kit following the manufacturer's protocol. The resulting pellet was resuspended in 1 mL of 1.2% SDS/PBS and sonicated for 3–4 s to obtain a homogenized solution. After heating it at 80–90 °C for 5 min, sample was cooled on ice. The sample was transferred to a 15-mL conical tube and diluted to 0.2% SDS with 5 mL of

PBS. Streptavidin-agarose beads (100 µL; pre-equilibrated with 3 × 200 µl PBS) were aliquoted into a 1.2-mL Bio-Spin column and washed three times with PBS (1 mL). Washed beads were suspended in 200 µL of PBS and added to the sample in the conical tube. The sample was rotated for 3 h at RT, and then centrifuged for 3 min at 1,400 × g. The beads were transferred into a Bio-Spin column, and the beads were washed with 10 mL of 0.2% SDS/PBS and then with 10 mL of PBS. The dried beads were transferred into a 1.7-mL eppendorf tube, to which 500 µL of a 6 M urea/PBS solution was added. In order to reduce the proteins, 25 µL of 200 mM Tris (2-carboxyethyl) phosphine (TCEP) was added to the sample, and it was heated at 65 °C for 15 min and then cooled to 37 °C. Alkylation of reduced proteins was accomplished by addition of 25 µL of 400 mM iodoacetamide (IAA) and incubation of the sample for 30 min at 37 °C with agitation in the dark. The sample was diluted with 950 µL of PBS, and then centrifuged at 1,400 × *g* for 2 min. The supernatant was discarded, and the beads were suspended in a premixed solution of 200 µL of 2 M urea/PBS, 2 µL of 100 mM $CaCl_2$ and 4 µL of 0.5 µg/ml trypsin. The reaction was allowed to proceed overnight at 37 °C with agitation, and then it was transferred to a 1.2-mL Bio-Spin column. After eluting the tryptic peptides into a low adhesion tube by centrifugation at 1,000 × *g* for 5 s, the beads were washed with 50 µL of water twice, and the eluent and water washings were combined in the same tube. Following the addition of 2 µL of formic acid (FA), the peptide sample was desalted using a ZipTip $C_{18}$ (see below).

**Preparation of Samples for Mass Spectrometry Analysis.** Ziptip ® C18 pipette tips (Millipore Sigma) were used to remove salts using the protocol from the manufacturer. MS grade water, acetonitrile (ACN) and trifluoroacetic acid (TFA) were used. Briefly, the dried samples were reconstituted with 13 µL of solution (R; 5:95, ACN:water, 0.1% TFA), vortexed and centrifuged (sample pH must be acidic at this point; TFA was added as necessary). The ZipTip was placed on a P10 pipettor set to 10 µL. The ZipTip was hydrated by aspirating solution 1 (50:50 ACN:water, 0.1% TFA) followed by solution 2 (0.1% TFA in water). Next, 10 µL of the sample was loaded by pipetting up and down 5-6 times. Next, the salts were removed by aspirating 10 µL of solution 2 (0.1% TFA in water) for a total of 4-5 washes (expel the wash into waste each time). Finally, the peptides were eluted by aspirating and expelling 10-20 µL of elution solution (60:40 ACN:water, 0.1% TFA) on ice 10 times (sample expelled into the same solution each time). Samples were stored at –20 °C until LC-MS/MS analysis.

**LC-MS/MS Analysis of Tryptic Digest.** Dionex Ultimate 3000 UHPLC system (NCS-3500RS pump and WPS-3000PL autosampler) was used for reverse phase chromatography. Five µL of sample (three biological replicates) was injected onto a home-packed analytical C18 reverse phase column with a 10 µm emission tip (75 µm × 200 mm, New Objective, Woburn, MA; Luna C18 5 µm particles, Phenomenex, Torrance, CA). Peptides were eluted with buffer A (0.1% FA in water) and buffer B (0.1% FA in acetonitrile) with the following gradient profile: 0-5 min for loading, 2% B, flowrate 1 µl/min; flowrate was decreased to 0.3 µl/min in 1 min; 6-21 min, 2%-10% B, flowrate 0.3 µl/min; 21-71 min, 10%-25% B, flowrate 0.3 µl/min; 71-100 min, 25%-40% B, flowrate 0.3 µl/min; 100-101 min, 40%-85% B, flowrate 0.3-1 µl/min; 101-106 min, 85% B flowrate 1 µl/min; 106-107 min, 85%-2% B, flowrate 1 µl/min; 107-115 min, 2% B, flowrate 1 µl/min. The coupled Thermo Elite mass spectrometer was operated in a Data-dependent acquisition mode, using Xcalibur v3.0 software. The global parameters were: ion source, NSI; ionization mode, positive; spray voltage, 2200 V; ion transfer tube temperature, 350 C; S-lens RF 50%, no sheath gas or auxiliary gas was used. For scanning mode, the orbitrap resolution was

30,000; scan range was 300-1800 *m/z*; AGC target was 1e6; maximum injection time was 100 ms; data type was profile. For MS2 mode, minimal signal threshold 5000; dynamic exclusion was enabled with a repeat count 1 and duration of 30 sec; the isolation window was 2 *m/z*; detector was orbitrap; top 10 ions were fragmented; activation type was HCD with a collision energy of 35%; orbitrap resolution was 15,000; AGC target was 5e4; maximum injection time was 100 ms.

MS raw files were processed using MaxQuant software (version 1.6.2; Max Planck Institute of Biochemistry). MS/MS-based peptide identification was carried out using the Andromeda search engine against the *B. subtilis* 168 UniProt database. For label-free protein quantification, the MaxLFQ algorithm was used as part of the MaxQuant package. The 'match between runs' option was enabled to maximize the number of quantification events across all replicates. Statistical analysis was performed in Perseus (version 1.6.1.1). Proteins identified only by site modification, reverse hits or contaminants were removed. Volcano plots were generated by performing a two-sample t-test with permutation-based statistics (FDR 0.01, S0 = 2).

**Antimicrobial Susceptibility Assay**

*B. subtilis* strain PY79 was streaked from the glycerol stock to LB-Agar plates and incubated at 37 °C for 14 to 16 h. Four colonies were inoculated into 5 mL of LB broth and grown to an $OD_{600}$ of 0.08-0.13. Cultures were diluted 100-fold (2.8 x $10^5$ CFU/mL). Solutions of antibiotics (0.125 – 64 μg/mL) in LB broth as two-fold dilutions were prepared. *Lower or higher concentrations were prepared as needed.* In 96-well plates, 100 μL diluted culture was added to 100 μL of serially diluted antibiotic in each well. Each compound was run in duplicate. After incubation at 37 °C for 18 to 20 h, plates were examined by eye. The lowest concentration that inhibited cell growth was recorded as the MIC.

**Fluorescence Microscopy**

Cells were grown in S750 minimal medium at 37 °C to an optical density ($OD_{600}$) of 0.3.(6) Cells were labeled with MEM-BODIPY and Click-MEM as described above, except that PBS was substituted with S750 minimal media to ensure cell viability. Microscopy was performed with a Nikon eclipse 80i microscope, equipped with a Plan Apo 100X Ph 3 objective (NA 1.4). Cells were mounted on 1% (wt/vol) agarose pads containing S750 minimal medium on object slides. Images were acquired with a Cool snap HQ2 camera (Photometrix) and were processed with Metamorph 7.7.9 software (Universal Imaging Corp.).

### Probe Synthesis and Characterization

#### Synthetic Procedures

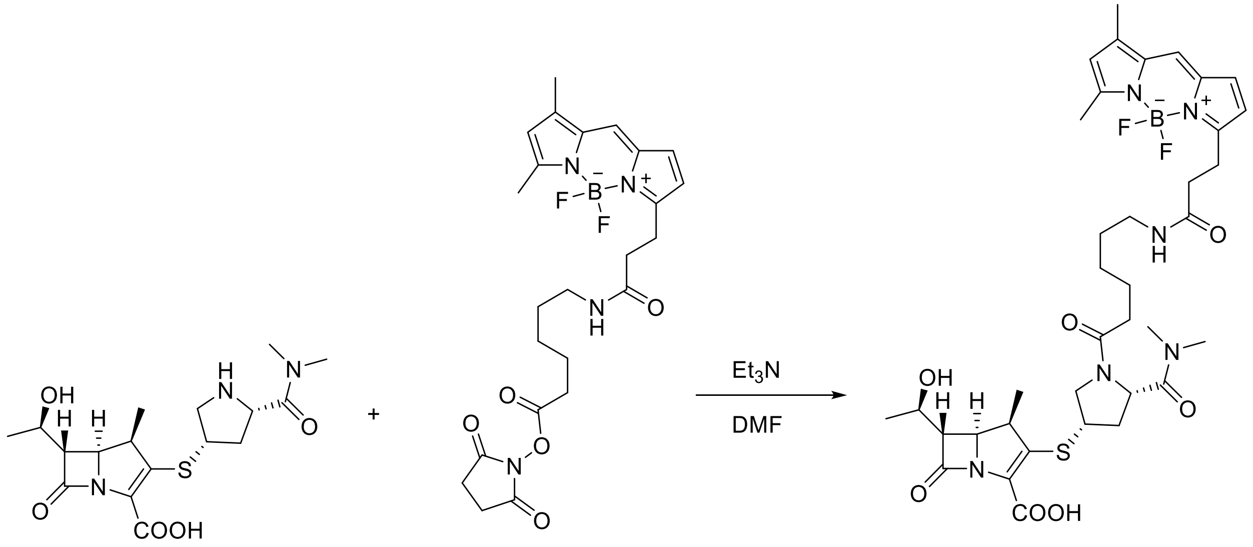


*MEM-BODIPY*

To a solution of BODIPY FL-X, SE (1.0 mg, 0.002 mmol, 1.0 equiv) in anhydrous DMF (0.1 mL) at room temperature was added meropenem (0.8 mg, 0.0022 mmol, 1.1 equiv) and TEA (0.0022 mmol, 0.3 µL, 1.1 equiv) in DMF (0.1 mL) and the mixture was stirred under Ar. Once the reaction was completed according to HPLC analysis, the solvent was evaporated and residue was purified by HPLC using a linear gradient of acetonitrile as indicated in the general methods section over 30 minutes on an Agilent Eclipse plus C18, 5 µm particle size, 4.6 × 100 mm at a flow rate of 2 mL/min (retention time = 12.3 min).

Reaction time of 5 days. Yield = not determine, only a portion of the reaction as fully purified (see example of purity in chromatogram below), HRMS-ESI: calc for $C_{37}H_{50}BF_2N_6O_7S^+$ $(M + H)^+$ 771.3517, found 771.3471. calc for $C_{37}H_{49}BF_2N_6NaO_7S^+$ $(M + Na)^+$ 793.3337, found 793.3287.

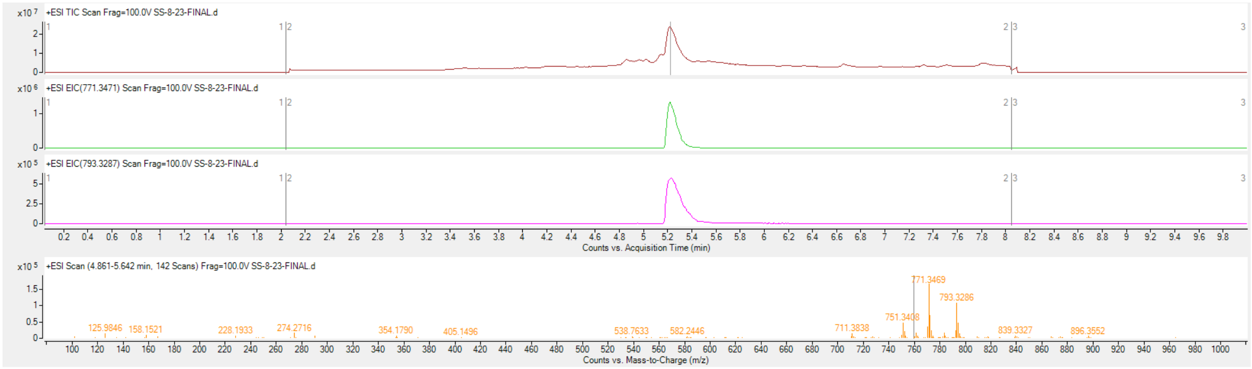

**Figure S14.** LC-MS analysis of MEM-BODIPY FL. The probe was dissolved in ~ 50 µL solvent A [$H_2O$, 0.1% formic acid (FA)] and analyzed on an UPLC-ESI-TOF MS instrumentation (Agilent, 6220; column: Agilent Eclipse plus C18, 5 µm, 100 × 4.6 mm). Gradient consisted of 5–95% B ($CH_3CN$, 0.1% FA) over 10 minutes. 100-1,000 Da mass range was detected. The compound was a single peak (retention time = 5.2 min), which contained protonated and sodium adduct of the MEM-BODIPY FL probe.

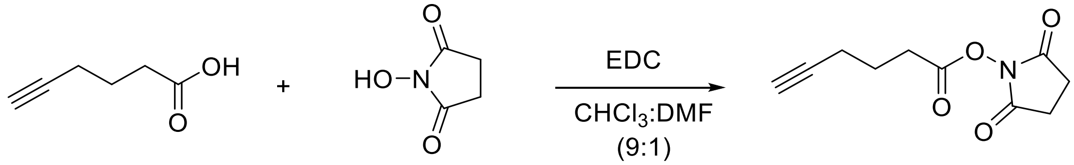


*N-Succinimidyl-5-hexynoate*

5-Hexynoic acid (1.0 g, 8.3 mmol) was dissolved in $CHCl_3$/DMF (5 mL, 9:1) and stirred. *N*-(3-dimethylaminopropyl)-*N'*-ethylcarbodiimide (EDC; 1.58 g, 8.3 mmol) and *N*-hydroxysuccinimide (0.96 g, 8.3 mmol) were added to the solution and the reaction stirred overnight under Ar. The reaction mixture was then diluted to 100 mL with dichloromethane and washed with 3 × 50 mL 0.1 N HCl, followed by 3 × 50 mL saturated $NaHCO_3$ and 1 × 50 mL saturated NaCl, dried over $MgSO_4$ and concentrated *in vacuuo* to yield 1.14 g yellow oil, which was taken to the next step without further purification. $R_f$ ~ 0.7 (3:7 EtOAc:$CH_2Cl_2$). Yield = 65%, $^1$H NMR (500 MHz, $CDCl_3$): δ 1.93-2.01 (quin, *J* = 7.2 Hz, 2H), 2.01 (t, *J* = 2.6 Hz, 1H), 2.32-2.36 (m, 2H), 2.75-2.78 (t, *J* = 7.2 Hz, 2H), 2.83 (br, 4H).

*Click-MEM*

To a solution of meropenem (3.8 mg, 0.001 mmol, 1.1 equiv) and cesium carbonate (3.2 mg, 0.01 mmol, 1.1 equiv) in anhydrous DMF (0.2 mL) on ice bath, was added *N*-succinimidyl-5-hexynoate (1.9 mg, 0.009 mmol, 1.0 equiv) in DMF (0.1 mL) and the mixture was stirred under Ar for 1 h until it reached room temperature. Next, the solvent was evaporated under vacuum and residue was purified by HPLC using a 0-20% gradient of acetonitrile over 20 min on an Agilent Eclipse plus C18, 5 µm particle size, 4.6 × 150 mm at a flow rate of 3 mL/min (retention time = 14.0 min). Yield = 35%, $^{1}$H NMR ($CD_3OD$, 500 MHz): δ 1.28 (d, J =7.3 Hz, 3H), 1.31 (d, *J* = 6.3 Hz, 3H), 1.71-1.79 (m, 2H), 1.82 (quin, *J* = 7.0 Hz, 2H), 2.25-2.31 (m, 3H), 2.46-2.59 (m, 2H), 2.75-2.81 (m, 1H), 2.97 (s, 3H), 3.19 (s, 3H), 3.29 (dd, *J* = 6.1, 2.55 Hz, 1H), 3.47 (t, J= 10.2 Hz, 1H), 3.56 (quin, J=7.3 Hz, 1H), 3.81-3.89 (m, 1H), 4.13 (quin, J = 6.5 Hz, 1H), 4.21-4.26 (m, 1H), 4.92 (t, J = 8.5 Hz, 1H). *$^{13}$C NMR could not be obtained due to rapid hydrolysis of the compound in organic solvents (soluble in methanol and DMSO, both of which caused hydrolysis over the course of the NMR experiment) and poor solubility in $D_2O$.* HRMS-ESI: calc for $C_{23}H_{32}N_3O_6S^+$ $(M + H)^+$ 478.2006, found 478.2031. calc for $C_{23}H_{31}N_3NaO_6S^+$ $(M + Na)^+$ 500.1826, found 500.1846.

**NMR spectra**

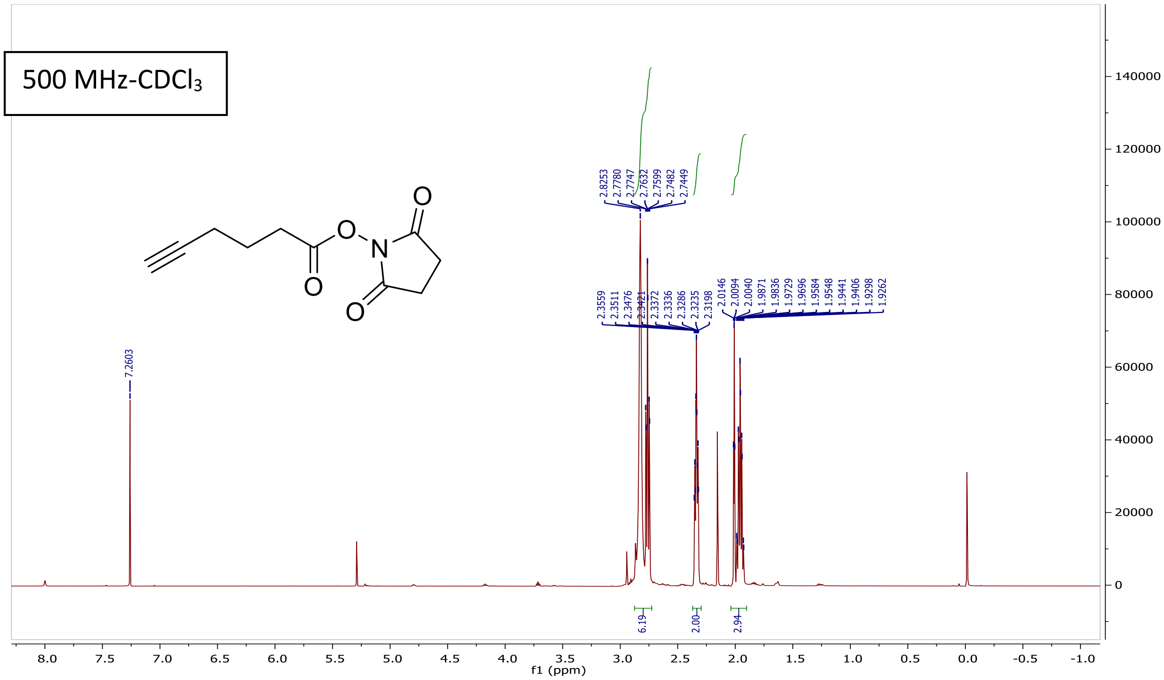


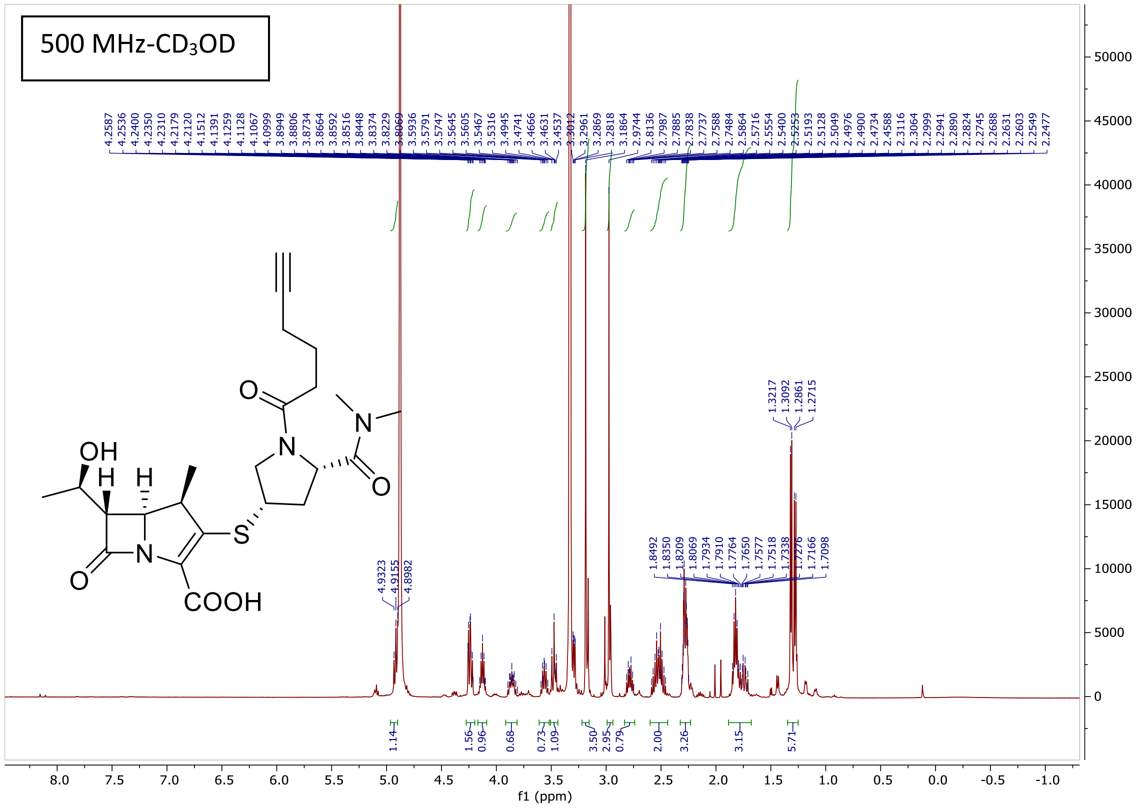
